# An interpretable open platform for sequence-based antibody developability prediction

**DOI:** 10.64898/2026.09.16.750421

**Authors:** Hoan N. Nguyen, Deepash Kothiwal, Yang Su, Minh Anh Kieu, Ruili Cao, Curtis Walton, Haisun Zhu, André A. R. Teixeira

**Affiliations:** Institute for Protein Innovation, Boston, MA 02115, USA

**Keywords:** antibody developability, machine learning, polyreactivity, protein language models, interpretable prediction

## Abstract

Antibody developability is increasingly predictable from sequence, yet software and trained models are rarely made available. We present DELPHI, open software for training developability predictors from labelled antibody assay data, together with ready-to-run, retrainable models. DELPHI compares 25 language-model and classifier combinations under CDR H3-cluster cross-validation that reduces sequence-similarity leakage, measures how performance changes with labelled training-set size, and reports residue-level model attributions. Applied to in-house polyreactivity and size-exclusion (SEC) data, it reaches mean AUC 0.959 and 0.933. Trained on those data alone, it transfers to a 246,293-antibody public library (AUC 0.950 with our deployed model) and ranks polyreactivity at a level similar to the best reported Ginkgo competition point estimate, without training on its data. Any laboratory can screen candidates before running assays, generate residue-level engineering hypotheses, and retrain DELPHI for a new assay.

## Introduction

Monoclonal antibodies are developed for both therapeutic and reagent applications, with more than 100 products approved and hundreds more in clinical development (1). By 2013, more than 1.4 million research antibodies were commercially available (2). A substantial fraction of candidates fail due to biophysical liabilities collectively termed “developability liabilities”, often discovered late in the pipeline at significant cost. Polyreactivity, i.e., the non-specific binding of an antibody to multiple unrelated proteins, is associated with accelerated clearance, immunogenicity, and diagnostic-assay interference (3–6). Aggregation and other non-monomeric failure modes compromise manufacturability and patient safety (7,8). Identifying these liabilities from sequence, early in discovery, can reduce later re-engineering (9,10).

Modern display technologies generate libraries of 10^6^–10^10^ diverse sequences, and next-generation sequencing (NGS) can recover hundreds of thousands of candidate binders per campaign, outpacing experimental developability screening (9,10). Sequence-based machine learning (ML) offers a route to close this gap through in silico pre-screening before costly assays. Protein language models (PLMs) capture features useful for antibody sequence analysis and developability prediction (11–15), and when combined with downstream classifiers or fine-tuned have shown promise for predicting polyreactivity (16–18), SEC monomer purity (13), and other developability properties using sequence or structural information (19–21). Yet most published approaches are not released as reusable, retrainable tools and interpret predictions at the level of features or sequence regions rather than individual residues, leaving experimental biologists without an integrated platform that combines multi-PLM benchmarking, cluster-aware cross-validation, training-set sizing, and per-residue attribution in one tool. Interpretable ML methods exist for biological sequence analysis (22) and for protein language model features (23), but integrated residue-level attribution remains uncommon in reusable antibody developability tools. Antibody developability prediction remains an open challenge: in the 2025 Ginkgo Developability Competition (24), the best PR-CHO submission achieved a Spearman |ρ| of 0.337.

To address these gaps, we built DELPHI (Deep End-to-end Learning Platform for antibody developability with High Interpretability), an open-source pipeline that lets a laboratory build, benchmark and deploy sequence-based developability classifiers from a single command line. DELPHI couples three capabilities. First, flexible model construction: five classifier architectures (XGBoost, Random Forest, CNN, a Transformer on language-model embeddings, and a dual-branch one-hot Transformer; Figs. S1–S3) across five antibody-specific PLMs plus one-hot, biophysical and k-mer features, giving 25 ready-to-benchmark combinations. Second, leakage-aware training: label curation (Fig. S4), class-imbalance correction, CDR H3-cluster-stratified cross-validation, and learning-curve analysis that characterizes performance across training-set sizes. Third, multi-resolution interpretability: SHAP (25) attribution for the tree-based models and Integrated Gradients (26) on a dual-branch one-hot Transformer that resolves attribution to individual residues across the full VH + VL + CDR H3 sequence. A user supplies labelled sequences and a configuration and obtains cross-validated predictions, interpretability outputs and a deployable checkpoint; models can be configured for binary biophysical readouts or run directly from the shipped IPI-trained PSR and SEC weights. Hyperparameters auto-scale to dataset size and class balance and remain fully configurable in YAML (Methods D, E).

Using DELPHI, we systematically benchmark all 25 model combinations under within-distribution cross-validation, cross-library transfer, and external clinical validation. PLM choice is a stronger performance determinant than classifier architecture, and framework-centric paired-antibody PLMs (AbLang2, IgBert) consistently lead PSR cross-library generalization. In DELPHI’s benchmarks, the more CDR H3-diverse IPI library showed stronger transfer despite being smaller, consistent with a contribution from training-set diversity: IPI-trained models transfer to a 246,293-antibody public library (AUC up to 0.950) (16) and, in zero-shot transfer, to three external clinical-stage cohorts with predictive transfer in the expected direction (Jain-panel AUC 0.73; Ginkgo PR-CHO |ρ| = 0.35, similar to the best reported competition point estimate). Multi-resolution interpretability and CDR3 loop in silico mutagenesis analysis (ΔP(Pass) relative to WT) identified a recurring CDR H3 electrostatic risk association (arginine and lysine enrichment, aspartate depletion, CDR3 length, net positive charge) across polyreactivity and SEC monomer purity failure. This is consistent with the heavy-chain charge determinants reported by Chen et al. (16) and shows a similar association with a second liability. VH and VL framework regions contribute model-attributed signal, while paired PLMs lead PSR cross-library transfer.

## Results

### Curated, CDR H3-diverse PSR and SEC training sets

The IPI dataset was generated from yeast display Fab libraries in which sequence diversity is confined to CDR H3 across eight VH and six VL germlines (Methods A). PSR-ELISA against four non-specific antigens (DNA, avidin, insulin, OVA/SMP) generated continuous polyreactivity scores for 7,837 antibodies; after denoising (Methods A; Fig. S4), 7,494 high-confidence binary-labeled antibodies were retained (∼79%/21% Pass/Fail). To address class imbalance, 3,771 high-polyreactivity Fail antibodies identified by NGS-based ssDNA selection were added, producing a near-balanced IPI PSR training set of 11,265 antibodies (53% Pass /47% Fail; Fig. 1b). The end-to-end DELPHI platform architecture is outlined in Fig. 1a. The IPI SEC dataset includes binary annotations for 5,045 antibodies based on size-exclusion chromatography profiles (∼64% Pass / 36% Fail), after retaining all Fail antibodies and selecting a subset of Pass antibodies using out-of-fold RF/XGBoost k-mer consensus scores as described in Methods B. For independent benchmarking, we used a published polyreactive dataset of 246,293 scFv (antibody set #1, or DS1) (16). CDR H3 diversity of IPI’s PSR dataset (7,263 clusters from 11,265 sequences) corresponds to ∼25-fold more clusters per sequence than the DS1 benchmark dataset (6,311 clusters from 246,293 sequences; Fig. 1c); this lower redundancy is associated with the transfer pattern described below. PSR pass rates also vary widely by VH germline (30–94% across germlines; Fig. 1d), showing an association between germline background and baseline polyreactivity pass rate. Biophysical-property profiling of the PSR and SEC training sets is provided in Figs. S5–S6.

**Fig. 1.**
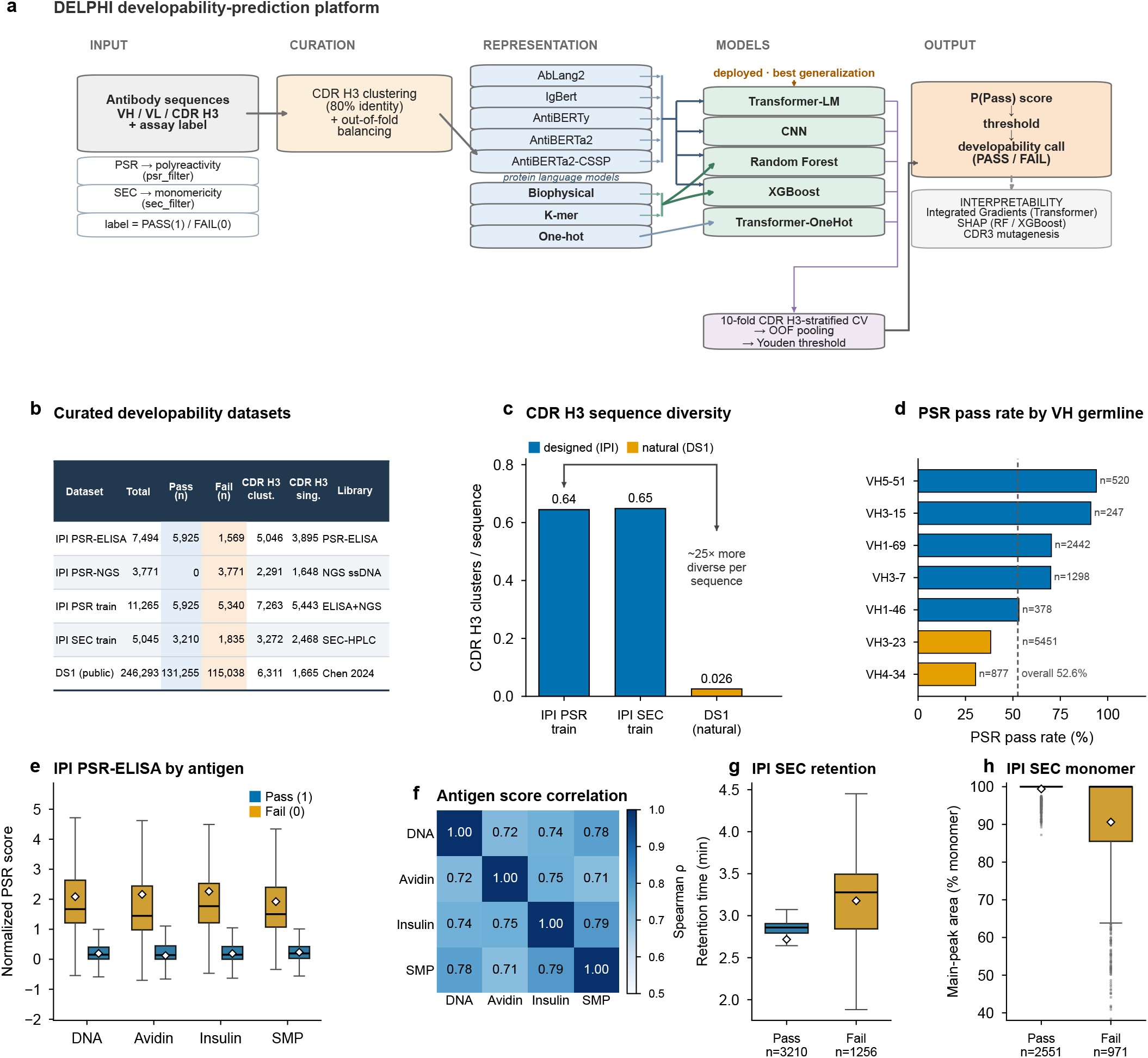
The DELPHI platform and the curated developability datasets. (a) DELPHI platform architecture: labeled antibody sequences (VH/VL/CDR H3) pass through curation (CDR H3 clustering at 80% identity, class balancing) into eight sequence representations (five antibody-specific PLMs plus biophysical, k-mer and one-hot encodings) and five classifier architectures, are pooled over 10-fold CDR H3-stratified cross-validation and thresholded at the Youden-J point, and return a P(Pass) score and a PASS/FAIL developability call; a parallel branch provides Integrated-Gradients and SHAP interpretability. (b) Summary of datasets used for training and benchmarking: total antibody count, Pass and Fail class sizes, CDR H3 clusters (80% Levenshtein identity), CDR H3 singletons, and originating library. The IPI PSR final training set (n = 11,265) combines PSR-ELISA-labeled antibodies (n = 7,494; 5,925 Pass / 1,569 Fail) and NGS-ssDNA-selected high-polyreactivity Fail antibodies (n = 3,771); the IPI SEC set (n = 5,045; 3,210 Pass / 1,835 Fail) was curated by SEC-HPLC; DS1 (n = 246,293) was used as published. (c) CDR H3 clusters per sequence for the designed IPI sets versus natural DS1; the designed library is ≈25-fold more diverse per sequence. (d) PSR pass rate by VH germline (germlines with n ≥ 100; dashed line, overall 52.6%), ranging from 30% (VH4-34) to 94% (VH5-51). (e) Normalized PSR-ELISA scores for Pass (blue) and Fail (orange) across the four assay antigens (DNA, avidin, insulin, OVA/SMP); box: IQR; line: median; diamond: mean; whiskers: 1.5× IQR. (f) Spearman correlation among the four antigen scores (ρ = 0.71–0.79), indicating a shared polyreactivity signal across assays. (g) SEC-HPLC main-peak retention time for available complete cases (n = 4,466; 3,210 Pass / 1,256 Fail) and (h) main-peak area (percent monomer) for available complete cases (n = 3,522; 2,551 Pass / 971 Fail); continuous chromatographic measurements were not uniformly available across historical assay records; box as in (e). Pass: low polyreactivity (PSR) or monomeric (SEC); Fail: high polyreactivity (PSR) or non-monomeric (SEC).

### Paired-chain PLMs generalize best across libraries

t-Distributed Stochastic Neighbor Embedding (t-SNE) (27) of the 11,265 IPI PSR trainset embeddings (ELISA + NGS) shows that all five PLMs organize antibodies primarily by VH germline: each germline occupies a distinct, contiguous region of the embedding space (Fig. 2a–e), and a 15-nearest-neighbour classifier recovers germline identity with ≥ 0.99 accuracy for every model (Fig. 2k). Re-colouring the identical coordinates by polyreactivity label (Fig. 2f–j) shows that Pass/Fail structure is also present but considerably weaker: a logistic classifier on the two t-SNE coordinates separates the classes with ROC-AUC 0.60–0.73, only modestly above the 0.5 chance line, and the ranking does not follow chain pairing (IgBert 0.73, AntiBERTa2-CSSP 0.70, AbLang2 0.69, AntiBERTy 0.62, AntiBERTa2 0.60; Fig. 2k).

**Fig. 2.**
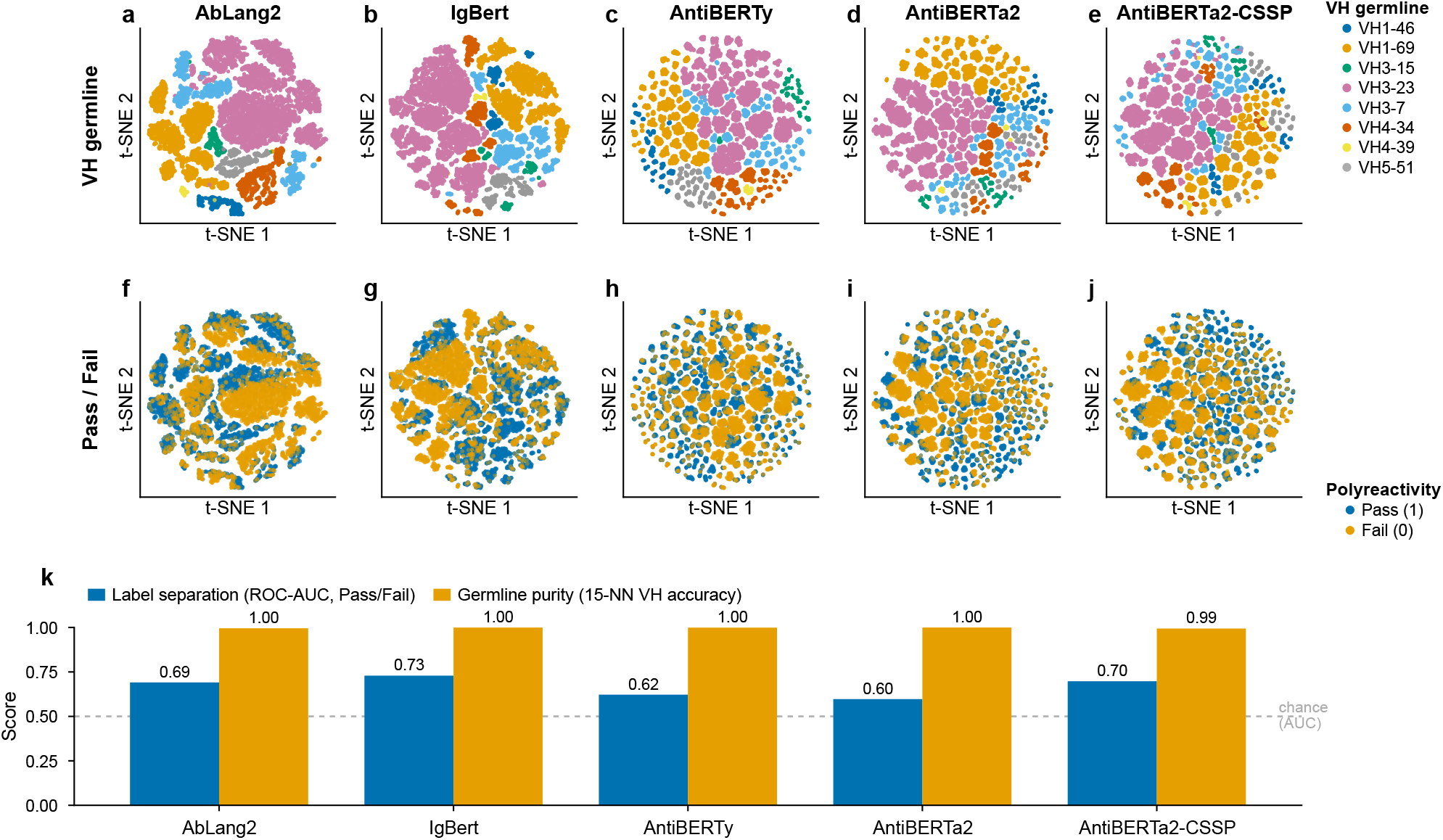
PLM embeddings organize by germline; developability-class structure is present but weak in every model. t-SNE of IPI PSR trainset embeddings (n = 11,265; ELISA + NGS) for AbLang2 (a, f), IgBert (b, g), AntiBERTy (c, h), AntiBERTa2 (d, i) and AntiBERTa2-CSSP (e, j), with identical coordinates colored by VH germline (top row, a–e) and by polyreactivity label (Pass/Fail; bottom row, f–j). (k) Quantification on the 2-D embedding per PLM: germline purity (15-nearest-neighbour VH-germline accuracy) and label separation (5-fold cross-validated ROC-AUC of a logistic model predicting Pass/Fail from the two coordinates). Because the t-SNE coordinates were fitted once on the complete cohort, these panels are descriptive visualizations rather than held-out predictive evaluations. Germline purity is near-perfect for all PLMs (0.99–1.00), whereas Pass/Fail separation is modest for every PLM (ROC-AUC 0.60–0.73; highest for IgBert at 0.73, lowest for AntiBERTa2 at 0.60) and does not follow chain pairing. t-SNE parameters: perplexity = 7, learning_rate = auto, max_iter = 1000, metric = euclidean, random_state = 42.

Germline framework identity thus dominates the embedding geometry, while the developability signal the downstream classifiers exploit is a finer structure layered on top. This raw-embedding separation is weak and does not track model architecture; the paired-chain advantage appears instead in cross-library transfer, where models that encode VH and VL jointly (IgBert, AbLang2) lead in both directions, as we quantify systematically across all 25 model combinations below.

### PLM choice outweighs architecture; transfer is asymmetric

Under 10-fold CDR H3-cluster-stratified cross-validation on the IPI PSR trainset (ELISA+NGS), all 20 PLM embedding-based architecture combinations achieve a mean within-distribution AUC of 0.959 ± 0.005 (range 0.946–0.967; Fig. 3d; Table S1), competitive with previously reported polyreactivity models (16,17). The best combination (XGBoost + IgBert, AUC 0.967) exceeds the 20-combination mean by 0.008 (≈1.4 cross-combination SD), a modest difference among closely performing models. Because the IPI library uses a restricted set of fixed framework designs, this within-library AUC is an optimistic estimate of generalization to new germlines: re-evaluating an XGBoost + AbLang2 classifier with entire VH germlines held out (leave-one-VH-germline-out) lowers its CDR H3-cluster pooled AUC of 0.965 to 0.903 (Methods E), and the cross-library analysis below provides a separate external transfer estimate. At the standard decision threshold (0.5), mean accuracy reaches 0.896 and mean F1 0.900 across all PLM combinations (Table S1). AUC is threshold-independent. DELPHI additionally computes a Youden-optimal operating threshold, which maximizes J = sensitivity + specificity − 1 and is embedded in each deployed checkpoint for inference; the Youden operating point is shown in the confusion matrices (Fig. 3e; Fig. S7). PLM choice is a stronger performance determinant than architecture: cross-architecture prediction agreement within the same PLM is high (Spearman ρ = 0.92–0.98; Fig. S8a). Traditional baselines (physicochemical descriptors, k-mer encodings; mean AUC 0.943) fall below the PLM range: the strongest baseline (XGBoost + biophysical) matches the weakest PLM combination (both AUC 0.946), and the best PLM combinations exceed every baseline. Transformer architecture with One-hot encoding (Fig. S3) performs competitively within the same library (AUC 0.961) but does not generalize across libraries (Fig. 4a); it is best suited to within-distribution screening and residue-level IG interpretation.

**Fig. 3.**
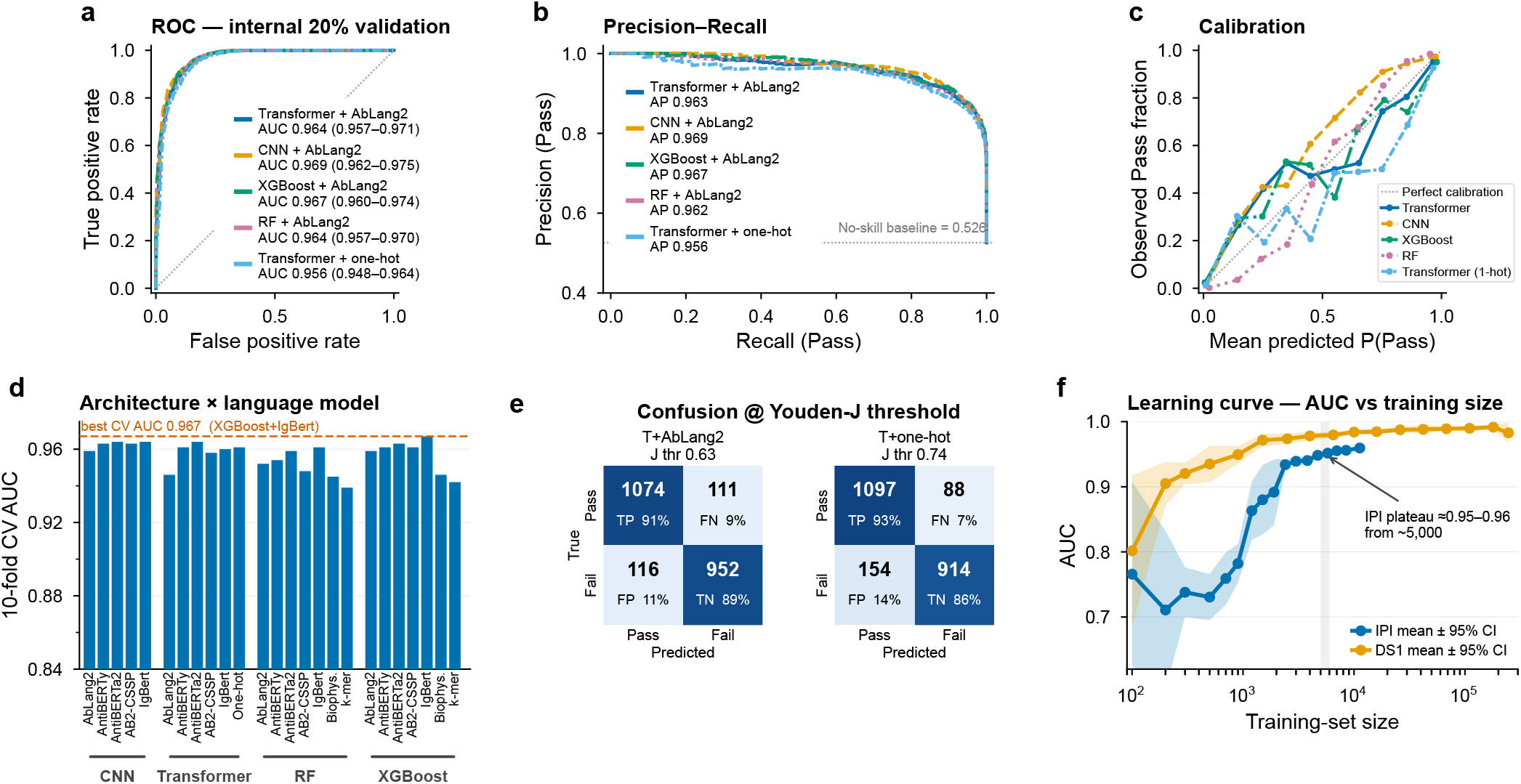
Within-distribution benchmarking on the IPI PSR data. (a) ROC curves on the held-out 20% validation split (n = 2,253; 1,185 Pass / 1,068 Fail) for five representative models (Transformer, CNN, XGBoost and Random Forest on AbLang2 embeddings, plus Transformer on one-hot); legend gives AUC with bootstrap 95% CI (2,000 resamples); per-antibody validation-split scores in Table S4. (b) Precision–recall curves for the same models (legend, average precision); the horizontal line marks the Pass prevalence in the validation set (no-skill baseline = 0.526). (c) Reliability (calibration) curves, 10 uniform bins; diagonal = perfect calibration. (d) 10-fold CDR H3-cluster-stratified cross-validated AUC for all 25 architecture × encoding combinations (20 PLM-based, plus Transformer + one-hot and RF/XGBoost biophysical and k-mer baselines) grouped by architecture; dashed line marks the best combination (XGBoost + IgBert, AUC 0.967). Full five-metric benchmark in Table S1. (e) Confusion matrices at the Youden-J operating point for Transformer + AbLang2 (threshold 0.63; sensitivity 0.91, specificity 0.89) and Transformer + one-hot (threshold 0.74; sensitivity 0.93, specificity 0.86); cells give counts and row-normalized percentages. (f) Learning curve: AUC versus training-set size for IPI (blue) and DS1 (orange) with points showing mean AUC ± 95% CI over repeated subsamples per size; gains slow markedly beyond ≈5,000 CDR H3-diverse antibodies (shaded band), where IPI plateaus near 0.95–0.96 and DS1 higher (≈0.98).

**Fig. 4.**
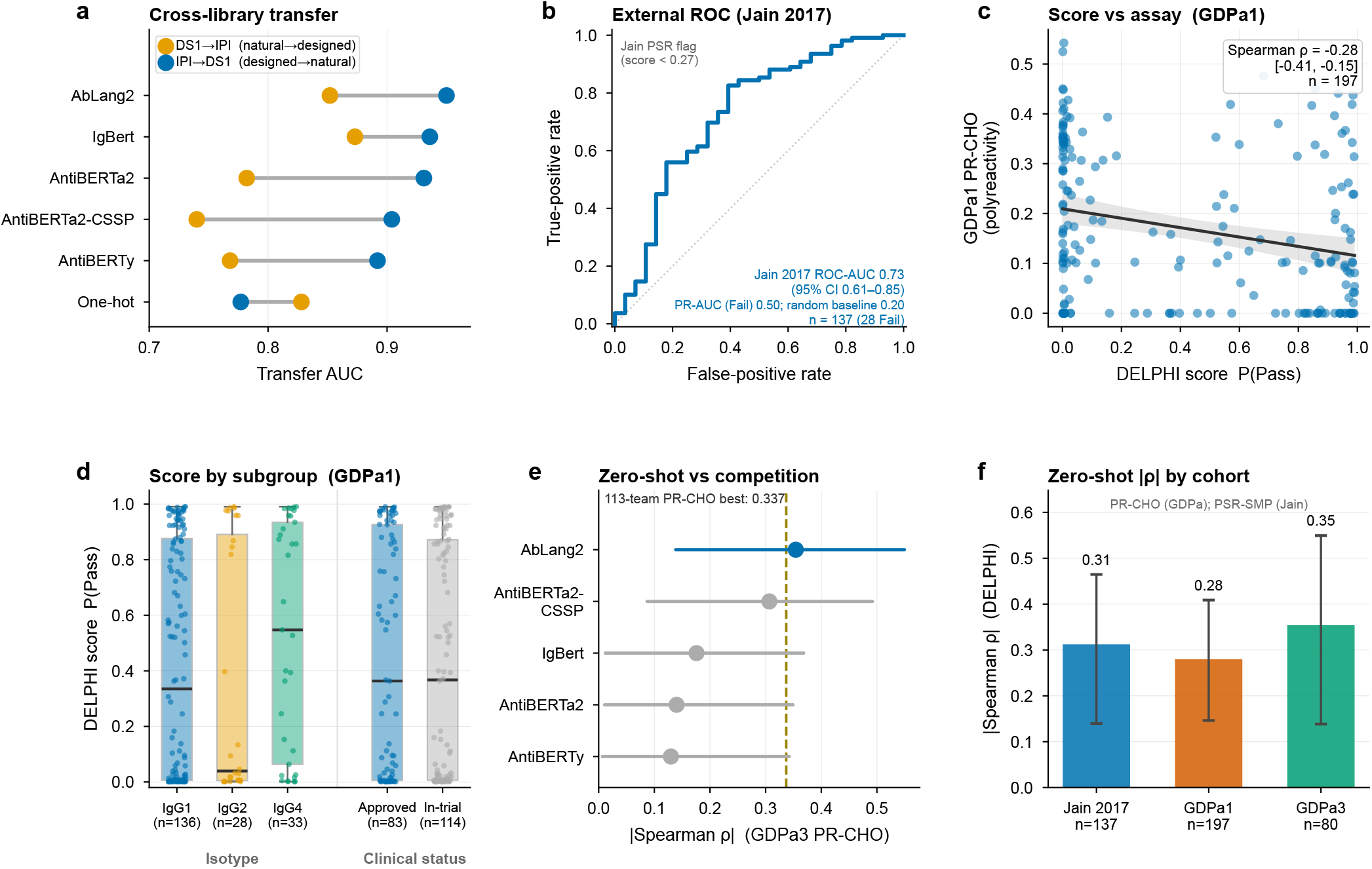
Zero-shot generalization to independent libraries and clinical-stage panels. All panels used the Transformer classifier architecture. (a) Cross-library transfer: per language model, transfer AUC for IPI→DS1 (designed→natural) and DS1→IPI (natural→designed) drawn as a dumbbell; AbLang2 reaches IPI→DS1 AUC 0.950 and IgBert DS1→IPI 0.883, whereas one-hot transfers worst (IPI→DS1 0.735). Full five-metric matrix in Table S2. (b) External zero-shot performance for the deployed Transformer + AbLang2 model on the Jain 2017 clinical-stage panel (n = 137; 28 Fail), binarized by the Jain PSR polyspecificity flag (score < 0.27): ROC-AUC 0.73 with bootstrap 95% CI and precision–recall AUC 0.50 for the rare Fail class (no-skill 0.20). The Ginkgo cohorts are evaluated by rank correlation (panels c, e, f), not a binary threshold. (c) DELPHI score versus the GDPa1 PR-CHO assay with OLS fit and 95% band; Spearman ρ = −0.28 (95% CI −0.41 to −0.15), negative as expected (higher P(Pass) = lower polyreactivity). (d) GDPa1 DELPHI score by IgG subtype and by clinical status (approved versus in-trial). (e) GDPa3 PR-CHO |Spearman ρ| per language model (point ± bootstrap 95% CI) against the best competition PR-CHO submission (|ρ| = 0.337, dashed line); AbLang2 reaches |ρ| = 0.35 (95% CI 0.14–0.55) in strict zero-shot transfer, similar to that best reported point estimate. (f) Best-model zero-shot |Spearman ρ| by cohort (Jain 2017, GDPa1, GDPa3). The full language-model × assay grid is in Fig. S9. Per-antibody prediction scores for the clinical-stage cohorts are in Table S5.

Cross-dataset transfer (Fig. 4a; Table S2) reveals an asymmetry consistent with differences in training-set CDR H3 diversity. DS1-trained Transformer models reach near-perfect internal AUC (mean 0.985, range 0.981–0.988) yet transfer poorly to IPI (DS1→IPI AUC 0.741–0.883; IgBert highest at 0.883), while IPI-trained models generalize strongly to DS1 (IPI→DS1 AUC 0.735–0.950; AbLang2 highest at 0.950) despite a 20-fold smaller dataset. At threshold = 0.5, the gap is also clear in F1: IPI→DS1 mean F1 0.831 versus DS1→IPI mean F1 0.743 (Table S2). Framework-centric paired-antibody PLMs (IgBert, AbLang2) lead on both transfer directions, consistent with paired VH+VL pretraining encoding germline and framework context that survives library changes, whereas positional one-hot encoding achieves competitive within-distribution performance and enables residue-level IG interpretation but shows substantially lower cross-library prediction agreement (Fig. S8b).

### Learning curves characterize performance across training-set sizes

DELPHI includes automated learning-curve analysis to characterize dataset-specific performance as training-set size increases. AUC was tracked as a function of training-set size for the deployed Transformer + AbLang2 model using CDR H3-cluster-stratified 80/20 splits (Fig. 3f; Methods E).

AUC is volatile below approximately 2,000 training antibodies (95% CI ∼0.6–0.9 at n ≤ 200 under repeated subsampling; Fig. 3f), then gains slow sharply and the IPI curve plateaus near 0.95–0.96 beyond ∼5,000 CDR H3-diverse antibodies (replicated mean ± 95% CI); DS1 plateaus higher, near ∼0.98. In these IPI and DS1 PSR analyses, approximately 5,000 CDR H3-diverse antibodies marked a practical performance plateau for the Transformer + AbLang2 model.

### Zero-shot transfer to clinical-stage panels

IPI-trained Transformer models were evaluated on three external clinical-stage cohorts: Jain 2017 (n = 137), GDPa1 (n = 197) (28), and GDPa3 (n = 80) across six polyreactivity readouts (Fig. 4b–f; Fig. S9). Across language models, zero-shot AUC on the two binarized Jain readouts ranges from 0.56 to 0.73 (Fig. S9a). On the Jain 2017 clinical-stage panel, binarized by the assay’s own polyspecificity flag (PSR score < 0.27, the 10%-worst-of-approved cutoff of Jain et al.), the deployed AbLang2 model reaches AUC 0.73 (95% CI 0.61–0.85; n = 137, 28 Fail), with precision–recall AUC 0.50 for the rare Fail class versus a 0.20 no-skill baseline (Fig. 4b). The Ginkgo cohorts (GDPa1, GDPa3) report PR-CHO on a different, bead-based scale (28) with no comparable Pass/Fail boundary, so we evaluate transfer there by rank correlation rather than a fixed threshold: Spearman ρ is negative in every cohort (ρ = −0.28 to −0.35 on PR-CHO; Fig. 4c, f; Fig. S9b), showing the expected inverse relationship between predicted P(Pass) and continuous polyreactivity score. On the blinded GDPa3 panel, DELPHI (AbLang2) transfers zero-shot to both polyreactivity readouts: |ρ| = 0.35 on PR-CHO (95% CI 0.14–0.55), similar to the best reported competition PR-CHO point estimate (|ρ| = 0.337; Fig. 4e), and |ρ| = 0.66 on PR-Ova (95% CI 0.50–0.79; Fig. S9). PR-Ova was not a competition scoring category and therefore provides an additional out-of-distribution transfer readout. The stronger PR-Ova correlation is consistent with ovalbumin also being one of the IPI PSR-ELISA antigens, whereas the PR-CHO readout (CHO-cell membrane proteins) is independent of the training antigens. DELPHI was trained exclusively on IPI PSR data; competition participants were provided GDPa1 as the development set, whereas GDPa3 was retained as the blinded test set (24).

### The framework extends to SEC monomer purity

To test whether DELPHI extends to another biophysical label, we applied it to SEC monomer purity failure, a distinct biophysical liability for which no large public training datasets are available. The IPI SEC trainset contained 5,045 antibodies, comparable to the approximately 5,000-antibody scale at which performance gains began to plateau for Transformer + AbLang2 in the IPI and DS1 PSR learning-curve analyses (Fig. 3f; Methods E). Across all 25 model combinations, AUC ranged from 0.8769 to 0.9596 (Table S3), with a mean AUC of 0.933 across the 20 PLM combinations, most of which clustered between 0.91 and 0.96. These results demonstrate that the DELPHI framework extends to a biophysical label beyond polyreactivity. XGBoost + biophysical achieves the highest SEC AUC (0.9596), with RF + k-mer (0.9581) and RF + biophysical (0.9573) close behind. These top results may partly reflect feature-space overlap with the k-mer-informed Pass-selection procedure (Methods B), rather than general model superiority; PLM embedding-based models therefore provide the more informative benchmark for this sequence-selected dataset. The SEC Pass/Fail labels are physical size-exclusion calls (monomer peak area and retention time; Methods B); the k-mer consensus step affected only downsampling of the Pass majority class, with all Fail antibodies retained. Because k-mer scores informed downsampling of the SEC Pass class, performance estimates may be modestly optimistic, particularly for k-mer-based models. The concordant direction of charge associations in the independently curated PSR dataset supports their robustness but does not eliminate this limitation. Pre-trained SEC model checkpoints are publicly available alongside the PSR checkpoints at https://doi.org/10.5281/zenodo.21823887, enabling immediate prediction of SEC monomer purity failure for new antibodies.

Predicted-score distributions visualize score separation (Fig. S10). Within the IPI PSR validation distribution, every architecture produces sharply bimodal scores, with Pass antibodies concentrated above P(Pass) 0.7 and Fail antibodies below 0.3 (Fig. S10a–e). On the full DS1 library, PLM-embedding models retain bimodal separation consistent with their within-distribution profiles, whereas the one-hot Transformer collapses toward P(Pass) ≈ 1 under distribution shift (Fig. S10f–j), the visual correlate of the cross-library transfer gap in Fig. 4a.

### A conserved CDR H3 charge signature in PSR and SEC

CDR H3 electrostatic imbalance is a prominent sequence-level correlate of both polyreactivity and SEC monomer purity failure (Methods C2). In the IPI PSR trainset (Fig. 5a–e; Fig. S5), PSR-Fail antibodies show marked enrichment of arginine (Fig. 5a), depletion of aspartate (Fig. 5b; Fig. S5b), and a positive shift in net nominal CDR H3 charge (Fail peak ∼0 to +2 versus Pass peak ∼−2 to 0; Fig. 5e). This sequence-derived charge shift supports the computational hypothesis that cationic CDR H3 composition contributes to non-specific interactions across structurally diverse PSR antigens. Longer CDR H3 loops are also associated with failure (Fig. 5d), although the contributions of conformational flexibility and solvent exposure were not measured directly. Tryptophan within antibodies carrying exactly one arginine (R count = 1) additionally associates with PSR failure (Fig. 5c), consistent with, but not establishing, aromatic contributions proposed for non-specific binding (29). Net CDR H3 charge shows the largest mean absolute standardized Pass/Fail separation across the three datasets (Cohen’s d = +2.18, +0.88 and +1.41 for IPI PSR, DS1 and IPI SEC; Fig. 5p), and the pass rate declines monotonically with net charge (from 0.80 at net charge 0 to 0.18 at +2; Fig. 5q). All five features are reproduced in DS1 (246,293 antibodies; Fig. 5f–j), consistent with the heavy-chain charge determinants reported for this same public library by Chen et al. (16); a similar association in the IPI SEC cohort is described below.

**Fig. 5.**
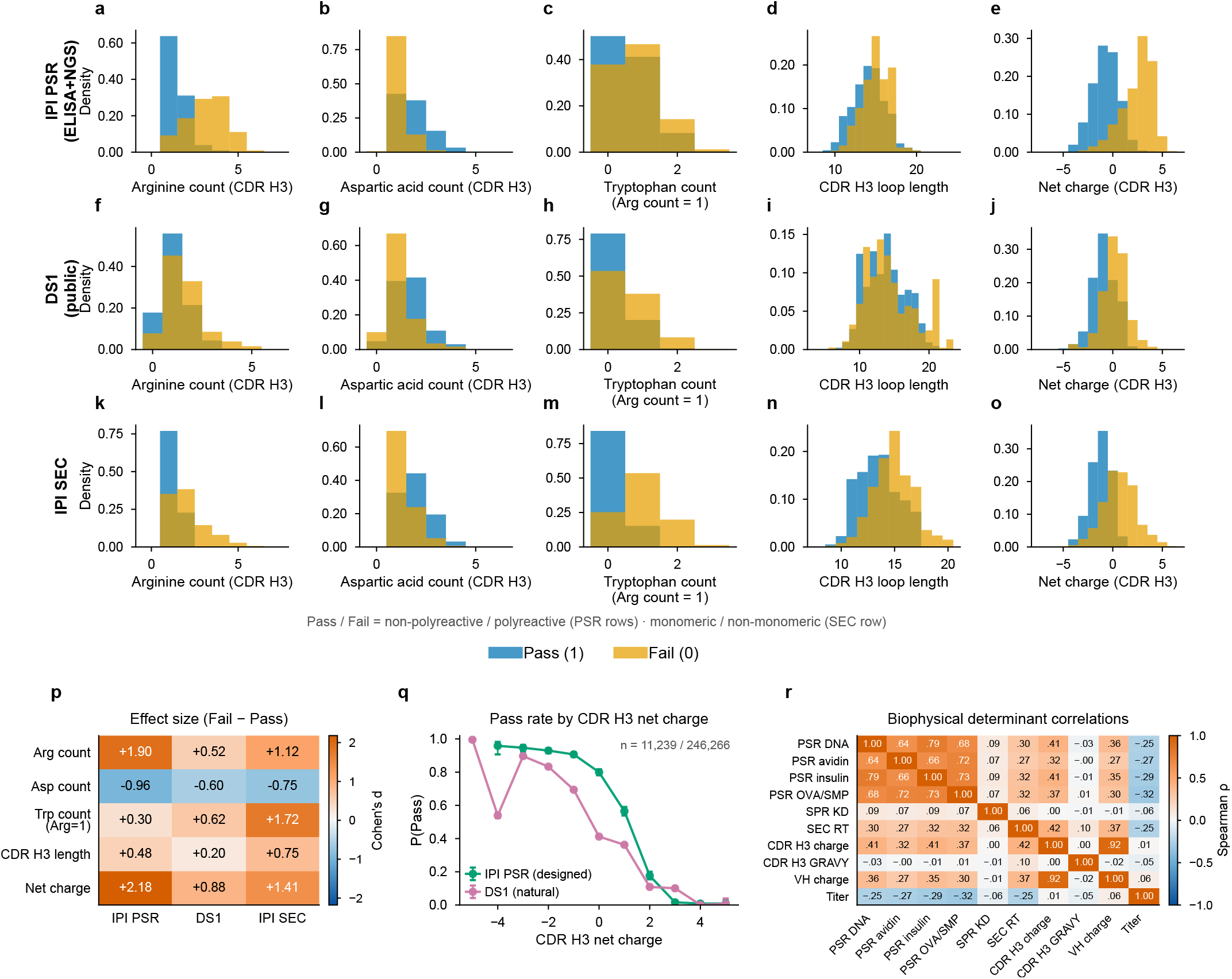
A conserved CDR H3 electrostatic failure signature across polyreactivity, SEC monomer purity failure, and independent libraries. (a–o) CDR H3 distributions of arginine (a, f, k), aspartate (b, g, l), tryptophan (c, h, m; arginine-count = 1 subset), loop length (d, i, n) and net charge (e, j, o) for Pass (blue) and Fail (orange). Row 1 (a–e): IPI PSR trainset (ELISA+NGS; n = 11,265); row 2 (f–j): DS1 (n = 246,293); row 3 (k–o): IPI SEC (n = 5,045). (p) Standardized Pass/Fail effect size (Cohen’s d, Fail − Pass) for each feature across the three datasets; net charge gives the largest mean absolute effect across the three datasets (d = +2.18 / +0.88 / +1.41 for IPI PSR / DS1 / IPI SEC). (q) Fraction Pass versus integer CDR H3 net charge for IPI PSR (green) and DS1 (purple); points are per-bin pass rate with Wilson 95% intervals, and only bins with n ≥ 20 are shown (11,239 IPI PSR and 246,266 DS1 antibodies represented); for IPI PSR, pass rate falls from 0.80 at net charge 0 to 0.18 at +2. (r) Available-case Spearman correlation matrix among biophysical determinants in the SEC cohort (PSR sub-assays, SEC retention time, CDR H3 and VH net charge and hydrophobicity, titer, and SPR equilibrium dissociation constant (KD, M)). Full 8-panel biophysical profiling is shown in Fig. S5 (IPI PSR) and Fig. S6 (IPI SEC).

Profiling of the IPI SEC trainset (Methods B) reveals similar CDR H3 composition and charge associations in non-monomeric antibodies (Fig. 5k–o; Fig. S6). SEC-Fail antibodies show arginine enrichment (Fig. 5k), aspartate depletion (Fig. 5l), extended loops (Fig. 5n), and net positive CDR H3 charge (Fig. 5o), mirroring all four PSR features. Within antibodies carrying exactly one arginine (R count = 1), tryptophan is modestly enriched in SEC-Fail (Fig. S6c), with greater class overlap than PSR at the distribution level, even though the trained models rank tryptophan as the strongest single feature for SEC failure (Fig. S11a, b). CDR H3 and whole-chain VH isoelectric points (pI) are both elevated in SEC-Fail (Fig. S6g, h); as CDR H3 is a primary contributor to overall variable domain charge, the whole-chain pI largely restates the CDR H3 charge signal. Together, these results identify a shared sequence-level signature in which arginine enrichment and anionic-residue depletion are associated with cationic CDR H3 loops and increased risk of both polyreactivity and SEC monomer-purity failure. This pattern is consistent with electrostatic promiscuity and reduced intermolecular repulsion as potential mechanisms, while fragmentation or structural instability may also contribute to SEC failure.

The convergence of polyreactivity and SEC monomer purity failure on the same CDR H3 physicochemical signature has direct engineering implications. Among the 5,045 antibodies measured in both assays, the two failure modes co-occur strongly (odds ratio 7.9; Fisher p < 10^−200^): PSR-Fail antibodies fail SEC in 71% of cases versus 24% for PSR-Pass, and CDR H3 net charge discriminates both labels within the same molecules (ROC-AUC 0.78 for PSR failure, 0.82 for SEC failure). Reducing net positive CDR H3 charge by targeting arginine enrichment and restoring anionic residues is predicted to reduce risk for both liabilities simultaneously, a sequence-based dual-liability engineering hypothesis. This physicochemical association is also reflected in ML attributions computed on the IPI data. TransformerOneHot Integrated Gradients localizes residue contributions to specific positions, whereas Random Forest and XGBoost SHAP analyses identify contributions from CDR H3 residue counts alongside higher-ranked aggregated descriptors. Both analyses show concordant Fail-associated effects of arginine enrichment and aspartate depletion (interpretability analyses, below).

### Attribution converges on a CDR H3 charge rule

We applied the same multi-resolution interpretability pipeline to PSR and SEC (Fig. 6).

**Fig. 6.**
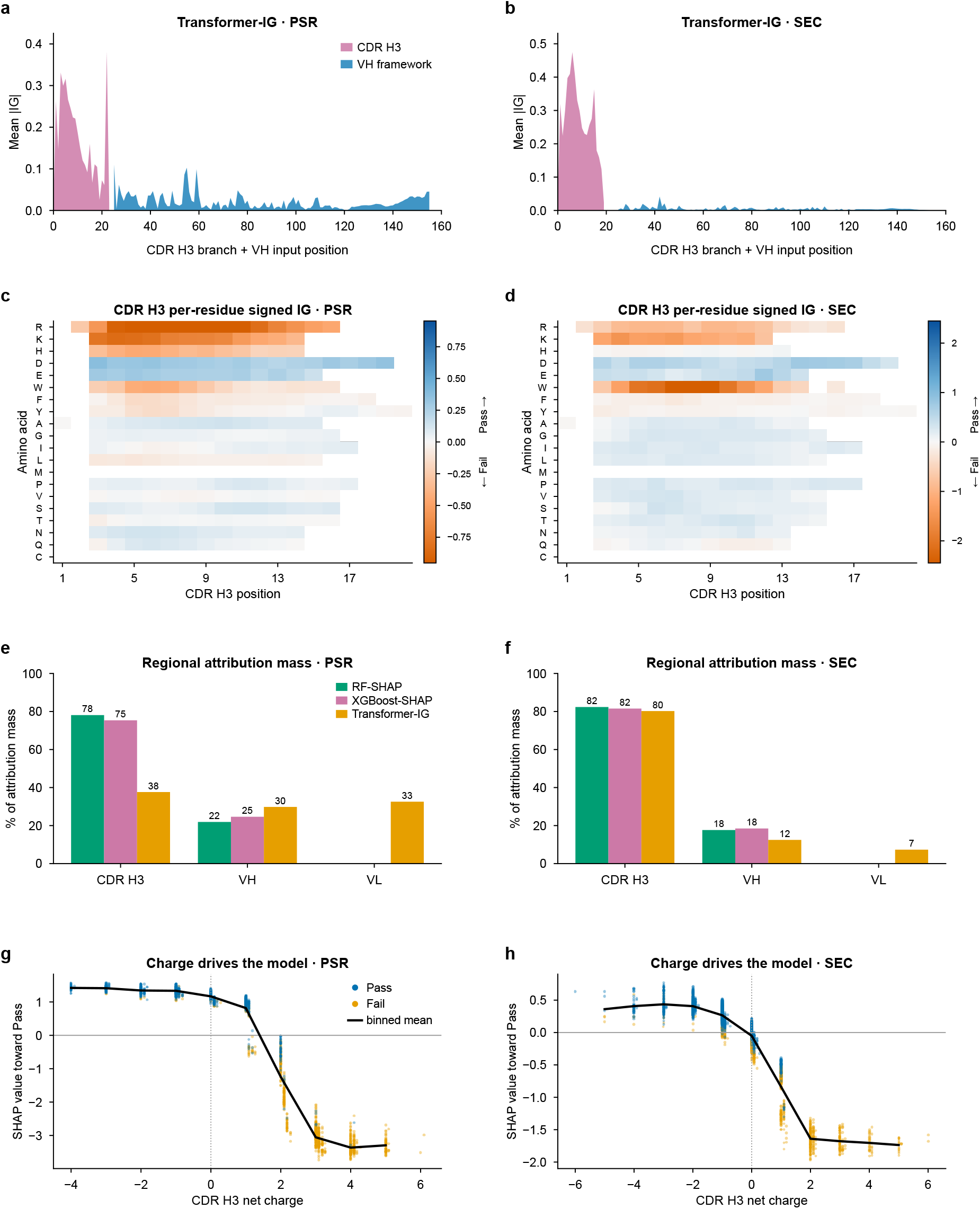
Multi-resolution interpretability reveals region- and residue-level determinants of antibody developability. (a, b) Mean absolute Integrated Gradients (|IG|) per input position for the CDR H3 branch followed by the VH portion of the VH+VL branch, for PSR (a) and SEC (b); mean per-position attribution is enriched in CDR H3. (c, d) Signed IG heatmaps for CDR H3 positions (amino acid × position), averaged among antibodies carrying each residue at each position, for PSR (c) and SEC (d); blue indicates attribution toward PASS and orange indicates attribution toward FAIL. Arginine (R), lysine (K), and tryptophan (W) are generally assigned toward FAIL, whereas aspartate (D) is generally assigned toward PASS. (e, f) Absolute attribution mass per region (CDR H3, VH, and VL), normalized separately within Transformer-IG, XGBoost-SHAP, and RF-SHAP, for PSR (e) and SEC (f). For PSR, Transformer-IG distributes attribution across CDR H3 (37.6%), VH (29.8%), and VL (32.6%), whereas the SHAP models concentrate more attribution in CDR H3; the SHAP models contain no VL features. For SEC, all three models assign more than 80% of absolute attribution mass to CDR H3. (g, h) XGBoost SHAP value for CDR H3 net charge versus net charge, coloured by Pass/Fail and overlaid with the binned mean, for PSR (g) and SEC (h); increasingly positive charge shifts the model output toward FAIL. Transformer IG uses all labelled antibodies (PSR n = 11,265; SEC n = 5,045), targets P(Pass), uses 200 integration steps, and uses a length-matched uniform amino-acid reference (1/20 at observed positions and zero only at true padding). XGBoost-SHAP panels use deterministic 3,000-antibody cohorts. Each signed IG heatmap cell reports the mean attribution among antibodies carrying that amino acid at that position; cells are shown only when at least 10 antibodies carry the indicated residue at that position, and uncertainty is not displayed.

Residue-level interpretation is performed using the TransformerOneHot model rather than PLM embedding-based models, despite the latter achieving higher cross-library transfer performance. The one-hot representation was selected because each input channel maps directly to an amino acid at a defined sequence position. Residue-resolved attribution through the frozen PLM pipelines was not implemented in this study. For the reported TransformerOneHot analyses, IG targeted the Pass class and used all labeled antibodies (PSR n = 11,265; SEC n = 5,045), 200 integration steps, and a length-matched uniform amino-acid reference (1/20 in each channel at observed positions; zero only at true padding). Median absolute convergence deltas were 4.47 × 10^−5^ for PSR and 4.19 × 10^−5^ for SEC. For interpretation, per-antibody signed IG reports the contribution assigned to each observed residue in that sequence. These attributions combine residue-position patterns learned by the model across the training data with sequence-context-dependent interactions in the individual antibody; IG does not separately decompose these components.

For PSR (Fig. 6e), attribution distributes broadly across all three regions, with CDR H3 (37.6%), VH (29.8%), and VL (32.6%) each carrying a comparable share of the total |IG| mass under the TransformerOneHot model; because VH and VL together span roughly ten-fold more residues than CDR H3, this reflects comparable total mass rather than per-residue importance, and mean per-residue CDR H3 attribution is substantially higher than mean VH attribution (Fig. 6a). For SEC (Fig. 6f), attribution concentrates strongly in CDR H3 (80.2%), with minor contributions from VH (12.5%) and VL (7.3%). Signed IG heatmaps at CDR H3 positions 1–20 (Fig. 6c, d) visualize position- and residue-specific model attributions: arginine (R), lysine (K), and tryptophan (W) are generally assigned toward FAIL at occupied CDR H3 positions, whereas aspartate (D) is generally assigned toward PASS, with broadly consistent directions between the separately trained assays. The dominant CDR H3 determinant differs by liability: net charge for polyreactivity and tryptophan content for monomer purity (Figs. S11–S12), on a shared, directionally conserved Arg-risk/Asp-protective background.

Agreement across three models with different feature spaces (Fig. 6e, f) provides cross-model computational consistency for this association. For PSR, TransformerOneHot-IG distributes attribution more broadly (CDR H3 37.6%, VH 29.8%, VL 32.6%), while SHAP models concentrate in CDR H3 (RF-SHAP 78.1%, XGBoost-SHAP 75.4%). RF/XGBoost use only CDR H3 + VH biophysical descriptors. TransformerOneHot was designed to capture long-range dependencies and inter-region interactions across the full VH + VL + CDR H3 sequence via dual-branch cross-attentio (Methods D), enabling attribution to any antibody region based on its contribution to the model prediction, a capability unavailable to biophysical descriptors and amino acid counts computed independently per region without positional or cross-region context. For PSR, this indicates a non-zero VL framework signal (33% of total |IG| mass, though far lower per residue than CDR H3); for SEC the VL signal is smaller (7%). This framework estimate comes solely from the TransformerOneHot-IG model and cannot be cross-checked by the SHAP models, which encode no VL features by construction; it should therefore be read as a single-method estimate. For SEC, all three models converge on CDR H3 (>80%), including TransformerOneHot-IG despite encoding VL; this convergence should be interpreted with caution as the SEC Pass majority class was downsampled using RF/XGBoost k-mer consensus scores (Methods B), which may strengthen CDR H3-associated signal.

For PSR (Fig. S12a, b), CDR3 net charge is the dominant protective feature and elevated CDR3 isoelectric point (high pI) the strongest risk-increasing feature across both RF and XGBoost, with arginine count (cdr3_R), tryptophan content (cdr3_W), and VH hydrophobicity ranking as secondary risk factors. For SEC (Fig. S11a, b), CDR3 tryptophan content is the top-ranked feature in both Random Forest and XGBoost, ahead of isoelectric point, net charge, and aromaticity; high tryptophan, high pI, and high net positive charge each drive failure, while low net charge is protective. In both assays, SHAP ranks aggregated biophysical group descriptors at the top; in the current RF/XGBoost implementation, SHAP resolves contributions from explicit CDR H3 residue-count features, including lysine and arginine counts, but these aggregate input features do not preserve the sequence positions responsible for each contribution. TransformerOneHot-IG additionally identifies lysine (K) as an explicit position-specific risk driver alongside arginine (R) (Fig. S12c for PSR; Fig. S11c for SEC), providing a position-specific view unavailable from aggregated biophysical descriptors that is directionally consistent with the aggregated SHAP descriptors. The Arg-risk/Asp-protective polarity is consistent across PSR and SEC. Plotting the XGBoost SHAP value for CDR H3 net charge against the net charge itself reveals a generally decreasing trend for both PSR and SEC (Fig. 6g, h), an explicit visualization of the learned model response.

Agreement between SHAP and TransformerOneHot-IG across three model families and two experimental readouts (Fig. 6e, f; Figs. S11–S12) provides cross-model computational consistency for a recurring CDR H3 electrostatic risk association. At individual antibody resolution, DELPHI generates per-antibody IG waterfall plots and CDR3 loop mutagenesis heatmaps that convert population-level signatures into residue-specific, testable engineering hypotheses (Fig. S13). As illustrated by Fail antibody TAB0015016 (Fig. S13b, d), CDR3_05_R, and CDR3_10_K carry the largest risk-increasing attributions across both PSR and SEC models. CDR3 loop in silico mutagenesis (Fig. S13e, f) shows that substituting the loop arginine and lysine residues with anionic (D, E) or neutral (A, G) residues increases ΔP(Pass) relative to WT. The in-silico mutagenesis results were directionally consistent with IG, SHAP, and biophysical-property analyses, providing cross-method computational corroboration.

## Discussion

DELPHI is an open-source, general-purpose framework for building and interpreting sequence-based developability classifiers across datasets. By benchmarking 25 model combinations across within-distribution cross-validation, cross-library transfer, and external clinical validation, it achieves strong within-distribution performance (mean AUC 0.959 for PSR and 0.933 for SEC across the 20 PLM combinations), transfers to 246,293 public PSR antibodies (AUC 0.950), shows predictive zero-shot transfer in the expected direction across three external clinical cohorts (Jain-panel AUC 0.73; on the blinded Ginkgo set, a point estimate similar to the best reported competition PR-CHO result), and provides multi-resolution model interpretation.

Table S6 compares DELPHI with the four most closely related methods. Chen et al. (16), Yu et al. (17) and Amini et al. (20) predict polyreactivity and related self-interaction assays, whereas Abeer et al. (13) predicts SEC monomer purity, collectively establishing that both liabilities are learnable from sequence. DELPHI extends this work in three practical directions: its trained PSR and SEC models are released as runnable, retrainable software; its learning curves quantify how predictive performance scales with labelled training-set size and provide assay-specific guidance for dataset design; and its attributions resolve predictions to individual residues.

Structure-based methods such as the Therapeutic Antibody Profiler model each variable domain and flag candidates by CDR length, surface hydrophobicity, and charge (30). For polyreactivity and monomer purity in these datasets, sequence provides substantial predictive signal: the models separate pass from fail within distribution, transfer across libraries (AUC 0.95), and recover a CDR H3 charge signature consistent with the electrostatics of non-specific binding (Fig. 5; Fig. S14b, c). These associations may help explain why sequence is predictive in these datasets and why transfer may vary across liabilities and libraries. Liabilities governed by the assembled fold rather than loop composition (a conformational aggregation motif or an interface epitope) may still require structure, and our fixed-framework library, varying only CDR H3, cannot test that case. Whether sequence suffices is therefore likely to be liability-specific. Where geometry drives failure, structure carries information that sequence does not, and combining the two is increasingly practical as antibody structure prediction improves.

Together, these findings define four practical contributions of DELPHI: quantitative guidance on data requirements, representation-aware model selection, assay-specific insight into framework dependence, and residue-level model interpretation.

First, learning curves translate performance benchmarking into a practical design rule: approximately 5,000 antibodies marked a performance plateau for Transformer + AbLang2 in the IPI and DS1 PSR analyses (Fig. 3f).

Second, PLM choice is a stronger determinant of predictive performance than classifier architecture: models sharing the same PLM embeddings produce highly correlated predicted scores across different architectures (Spearman ρ = 0.92–0.98; Fig. S8a). Framework-centric paired-antibody PLMs (AbLang2, IgBert) showed clear advantage for cross-library transfer in both directions, consistent with paired VH+VL pretraining encoding germline and framework context that survives library changes. Tree-based models (Random Forest, XGBoost) remain competitive classifiers and can serve as practical pre-screening tools for rapid large-library filtering.

Third, the value of paired-chain context is assay-dependent rather than universal. The paired antibody PLMs AbLang2 and IgBert lead for PSR and cross-library transfer, consistent with germline-dependent pass rates and germline-separated embeddings, whereas SEC shows near parity between paired and unpaired PLMs (Fig. 2; Fig. 4a; Table S3). The framework attribution from TransformerOneHot-IG is therefore supportive rather than definitive: it is distributed over many framework residues, arises from the weakest-transfer model, and cannot be independently tested with SHAP models that omit VL features (Fig. 6a, b).

Fourth, physicochemical profiling, TransformerOneHot-IG, RF-SHAP, and XGBoost-SHAP show consistent directions of association across PSR and SEC: cationic enrichment is associated with greater risk, whereas anionic residues are associated with lower risk (Figs. 5 and 6). Its recurrence in the molecule-level cross-assay test and CDR H3 loop mutagenesis provides cross-method computational corroboration of a dual-liability engineering hypothesis (Figs. S13, S14b, c).

Current IPI-trained models reflect the constraints of their training data: PSR labels derived from ELISA and NGS-based ssDNA selection (Methods A) may not capture all modes of polyreactivity, while SEC calls based on monomer peak area and retention time may not capture all non-monomeric failure mechanisms, the IPI germline space is restricted (8 VH, 6 VL germlines) with diversity confined to CDR H3, and whether predictions generalize to antibodies outside the IPI germline composition, including rare germlines and diverse framework scaffolds, remains to be established; consistent with this, holding entire VH germlines out at training lowers the within-library AUC from 0.965 to 0.903 (Fig. S14a). The PSR Fail class is drawn partly from a distinct NGS-ssDNA selection rather than the ELISA screen used for the Pass class, so the within-distribution PSR AUC is best interpreted as an upper estimate. Separately, IPI-trained models achieved cross-library transfer to DS1 with AUCs of 0.735–0.950. Local PLM fine-tuning has also improved identification of low-polyreactivity variants within an optimization campaign, with little change in performance on an external clinical panel (31).

Moving forward, DELPHI models for PSR and SEC, together with future models for viscosity, thermal stability, or immunogenicity, could be combined into a joint scoring function for multi-objective optimization within directed evolution, Gibbs sampling, or reinforcement-learning frameworks (32,33). Such an approach could enable simultaneous optimization of multiple developability properties, a multi-objective engineering goal (34) that depends on accurate, accessible property predictors.

Extending to continuous regression will enable quantitative score prediction beyond binary classification. Future implementations could extend residue-level attribution to PLM-based models, bridging the current gap between interpretability and cross-library performance. Adapting PLMs to laboratory-specific antibody sequences without assay labels may adapt representations to specific germline compositions and liability distributions, an approach recently shown to improve developability prediction on internal datasets (20) that DELPHI complements with open weights, multi-PLM benchmarking, and residue-level interpretability. Multi-task learning across PSR and SEC could model co-dependencies between liabilities. For the two liabilities and the germline-restricted libraries examined here, developability is partly predictable from sequence; whether this extends to broader scaffolds and assays remains open. DELPHI extends prior work in antibody developability prediction (9,10,19) with open, interpretable models that any laboratory can build from its own data and run or retrain from a single command.

## Materials and Methods

### A IPI PSR-ELISA assay, dataset curation, and denoising

Yeast display library construction. Fab libraries were constructed with sequence diversity confined exclusively to CDR H3 across eight VH germlines (VH1-69, VH3-23, VH3-7, VH4-34, VH5-51, VH3-15, VH1-46, VH4-39) and six VL germlines (κ1-39, κ3-15, κ3-20, κ4-1, and the lambda germlines VL1-51 and VL2-14); all framework regions were held fixed. Per-germline pass rates for the germlines with n ≥ 100 (seven VH, five VL) are shown in Fig. 1d. Library generation and yeast surface display protocols followed Kothiwal et al. (35).

PSR-ELISA protocol. Polyspecificity was measured by ELISA against four non-specific antigens: double-stranded bovine DNA (1 μg/mL in 1x PBS), ovalbumin (OVA; 5 μg/mL), insulin (5 μg/mL), and avidin (10 μg/mL). The OVA antigen replaced a soluble membrane protein preparation (SMP) used in earlier batches; scores from the two reagents are pooled as a single OVA/SMP polyreactivity readout. Plates were coated overnight at room temperature, blocked for 1 h, and incubated sequentially with primary antibody (1 h), secondary antibody (1:5,000 dilution, 1 h), and substrate (30 min). Reactions were quenched with 5N NaOH and absorbance read at 405 nm on a SpectraMax plate reader.

Dataset assembly. The IPI PSR dataset was assembled from libraries screened by PSR-ELISA, yielding four continuous polyreactivity scores per antibody. Antibodies with missing values in any of the four scores were excluded, producing a starting set of 7,837 antibodies with complete profiles. Binary Pass (1; non-polyreactive) and Fail (0; polyreactive) labels were assigned using an internal annotation method that integrates all four continuous ELISA scores into a single binary call; these labels serve as ground truth for all downstream ML analyses.

Label quality control. Raw PSR-ELISA score distributions exhibited substantial overlap near the decision boundary, reflecting genuine biological heterogeneity and assay measurement variability. To remove low-confidence label assignments we applied a two-stage quality control procedure. In the primary stage, a Random Forest (RF) classifier (36) was trained on the internally annotated binary labels using all four continuous ELISA scores as input features. Antibodies whose RF-predicted class disagreed with their assigned label were flagged as low-confidence, indicating that their score profile was inconsistent with the multivariate class boundary learned from the full dataset. In the confirmatory stage, SVM-based decision boundary modeling on continuous DNA and OVA/SMP scores, combined with t-SNE visualization of all four ELISA scores (perplexity=30, 1,000 iterations), verified that RF-flagged antibodies were concentrated in the ambiguous boundary region between Pass and Fail clusters (Fig. S4a–c), confirming that removed cases represented genuine label uncertainty rather than systematic misclassification. This procedure removed 343 low-confidence antibodies, yielding 7,494 high-confidence labels (5,925 Pass, 1,569 Fail; ∼79%/21%) with markedly improved class separation.

Class balancing. The ELISA-only set was substantially imbalanced (∼4:1 Pass:Fail), risking bias toward the majority class. To address this, 3,771 high-polyreactivity Fail antibodies independently identified by NGS-based ssDNA selection (read count >30) were added, yielding a near-balanced final IPI PSR training set of 11,265 antibodies (5,925 Pass, 5,340 Fail; ∼53%/47%). For cross-dataset benchmarking, the publicly available DS1 library was used without filtering, comprising 246,293 scFv antibodies with PSR labels (131,255 Pass, 115,038 Fail).

Dataset characteristics and consequences for generalization. Two structural properties of the IPI dataset have direct consequences for model generalization. First, germline composition shapes polyreactivity baseline: within the IPI PSR training set, VH germline-specific pass rates vary substantially: VH3-15 and VH5-51 show high polyreactivity tolerance (>88%) whereas VH1-46 shows lower tolerance (53%), consistent with germline-encoded surface charge biases that predispose certain frameworks toward non-specific binding (4). Second, CDR H3 sequence diversity differs dramatically between datasets: IPI yields 7,263 clusters from 11,265 sequences (∼48% singletons), whereas DS1 produces only 6,311 clusters from 246,293 sequences (<1% singletons), a ∼25-fold difference in relative CDR H3 diversity. This structural difference means IPI’s CDR H3-cluster cross-validation evaluates each model largely on CDR H3 sequences absent from its training folds.

CDR H3 sequence clustering. CDR H3 sequences were clustered using a greedy Levenshtein distance algorithm at 80% sequence identity. For two sequences of lengths L_1_ and L_2_, similarity is the Levenshtein ratio, (L_1_ + L_2_ − d) / (L_1_ + L_2_), where d is the indel edit distance (insertions and deletions, a substitution counting as two edits), computed with Levenshtein.ratio from the Levenshtein Python package. A sequence was assigned to an existing cluster if its Levenshtein ratio to the cluster seed exceeded 0.80; otherwise it seeded a new cluster. This keeps sequences that share a cluster seed (>80% identity to that seed) together across training and validation folds, reducing optimistic bias from sequence similarity leakage; because clustering is greedy by seed, a small fraction of >80%-identical pairs assigned to different seeds can still fall on opposite sides of a split.

Biophysical feature computation. CDR H3 net charge, isoelectric point (pI), and amino acid composition were computed using the Biopython ProteinAnalysis module (Biopython 1.87). Net charge at pH 7.4 was calculated using standard pKa values for ionizable residues (Arg, Lys, His, Asp, Glu, Cys, Tyr, N-terminus, C-terminus); pI was determined as the pH at which net charge equals zero using the Henderson–Hasselbalch equation.

### B IPI SEC-HPLC assay

SEC-HPLC protocol. Antibodies were transiently expressed in Expi293 cells and purified by Protein A affinity chromatography. Purified protein was diluted to 0.5 mg/mL in PBS and injected onto a TSKgel SuperSW mAb HTP (4 μm) HPLC column (Tosoh Bioscience) on an Agilent 1260 HPLC system. Separation was performed isocratically in PBS at a flow rate of 0.5 mL/min over a 6-minute run. Monomer peak identity was confirmed by retention time (2.677–3.251 min, calibrated against rabbit IgG standards). Antibodies were classified as Pass (1; monomeric; monomer peak area ≥ 90%, peak height ≥ 5 mAU) or Fail (0; aggregated or fragmented; monomer peak area < 90%).

Dataset denoising and rebalancing. The raw SEC dataset, compiled from multiple assays conducted over several years, comprised 7,019 antibody candidates. It exhibited a highly imbalanced class distribution (5,184 PASS / 1,835 FAIL, ≈74:26 ratio) along with substantial noise. The PASS class was further expected to contain borderline or mislabelled entries arising from assay variability inherent to analytical SEC. To address both imbalance and label noise, a combined diversity-confidence downsampling strategy was applied to the PASS class while all 1,835 FAIL samples were retained in full.

PASS candidates were first clustered by CDR H3 Levenshtein similarity (threshold = 0.8) to identify one representative per cluster, eliminating clonally redundant sequences and maximising CDR3 diversity in the selected subset. Concurrently, Random Forest and XGBoost (37) classifiers trained on 1–3-mer amino acid frequency features were used to score each PASS candidate via out-of-fold prediction (5-fold cross-validation, ROC-AUC optimized by grid search), ensuring that confidence estimates were unbiased by in-sample memorization. Candidates confirmed by both models at a consensus probability threshold of ≥ 0.6 were considered label reliable. Final selection prioritized candidates satisfying both criteria (cluster representative and consensus-confirmed), reducing the original 74:26 class imbalance to a more tractable 64:36 ratio and yielding a curated training set of 5,045 antibodies (3,210 PASS / 1,835 FAIL) for model training (Fig. 1). The final cohort contains complete curated binary SEC labels for all 5,045 antibodies. Continuous chromatographic measurements were not uniformly available across historical assay records collected over multiple years; analyses of retention time and main-peak area therefore used available complete cases without imputation (Fig. 1g,h).

### C Sequence encoding and protein language model embeddings

Embeddings and feature encodings are generated automatically on first use and cached to disk by antibody BARCODE identifier ({input_file}.{lm}.emb.csv); subsequent runs load the cached file directly, eliminating redundant computation across downstream classifiers.

#### C0. Input data requirements

DELPHI accepts CSV (.csv) or Excel (.xlsx, .xls) files. The minimum required columns differ by use case:

##### Training and cross-validation

| Column | Description |
|---|---|
| BARCODE | Unique antibody identifier |
| HSEQ | VH amino acid sequence |
| LSEQ | VL amino acid sequence |
| CDR3 | HCDR3 amino acid sequence |
| TARGET | Binary label column (e.g. psr_filter, sec_filter, spr_anno); Pass = 1, Fail = 0 |

Prediction on unseen antibodies:

| Column | Description |
|---|---|
| HSEQ | VH amino acid sequence |
| LSEQ | VL amino acid sequence |
| CDR3 | HCDR3 amino acid sequence |

BARCODE is optional for prediction; if absent, integer indices are assigned automatically. All sequence columns must contain single-letter amino acid strings. Rows with missing values in any required column are excluded prior to embedding generation or model training.

#### C1. Protein language model embeddings

Five antibody-specific protein language models (PLMs) were used to generate fixed-length per-antibody embedding vectors. All Transformer-based embeddings were computed using mixed-precision inference on GPU where available, with CPU fallback, and processed in batches of 64–128 sequences with padding and truncation to a maximum of 512 tokens.

##### AbLang2

(12) (480-dimensional). Based on the ESM-2 Transformer architecture (38), pre-trained on 35.6 million unpaired and 1.26 million paired VH/VL sequences from the Observed Antibody Space (OAS) database (39,40). Paired VH+VL embeddings were generated using ablang2.pretrained(“ablang2-paired”) with mode=‘seqcoding’, returning a single fixed-length vector per antibody pair.

##### AntiBERTy

(11) (512-dimensional). A BERT-based Transformer pre-trained on 558 million VH/VL sequences from OAS. Paired embeddings were generated by concatenating heavy and light chain sequences with a spacer token, applying spaced tokenization, and mean-pooling over non-padding token hidden states from the final encoder layer.

##### AntiBERTa2

(41) (1,024-dimensional). A RoFormer architecture pre-trained on 779.4 million unpaired heavy and light chain sequences from OAS. Per-antibody embeddings were generated by mean-pooling over non-padding token hidden states from the final encoder layer.

##### AntiBERTa2-CSSP

(41) (1,024-dimensional). A structurally supervised variant of AntiBERTa2, additionally pre-trained on 1,554 human antibody crystal structures using a contrastive sequence-structure pre-training (CSSP) objective that incorporates paired sequence–structure information. Embeddings were generated identically to AntiBERTa2.

##### IgBert

(14) (1,024-dimensional). A ProtBert-backbone (42) BERT model fine-tuned on more than 2 million paired VH/VL sequences from OAS. Paired sequences were encoded as [CLS] VH tokens [SEP] VL tokens [SEP] with amino acids space-separated. Embeddings were generated by mean-pooling over all non-padding token hidden states including [CLS] and [SEP] tokens, following the procedure recommended by the original authors.

#### C2. Biophysical feature extraction

CDR H3 and VH biophysical features were computed for all datasets (IPI PSR, IPI SEC, DS1) using two custom Python scripts available in the DELPHI repository: *utils/liabilities*.*py*, which implements all feature extraction functions, and *utils/Figure2_physicochemical*.*py*, which applies these functions to generate Fig. 5, Fig. S5, and Fig. S6. For DS1, heavy-chain CDR3 was annotated from the VH sequence with ANARCI (IMGT scheme, positions 105-117) prior to feature computation (utils/download_ds1_dataset.py). Per-residue counts of arginine (R), aspartate (D), glutamate (E), and tryptophan (W) were computed as integer frequencies over the CDR H3 sequence (*annotate_liabilities_2()*, cdr3_col=‘CDR3’). Net nominal CDR H3 charge was computed as (R + K) − (D + E) (*simple_charge()*). CDR H3 loop length was the number of residues in the CDR3 sequence; antibodies with CDR3 length ≥ 25 were excluded as likely sequencing artefacts. CDR H3 and VH isoelectric points (pI) were computed using *Bio*.*SeqUtils*.*IsoelectricPoint*. Additional per-sequence features including charge at pH 7.4, GRAVY hydrophobicity, aromaticity, and instability index were computed using Biopython’s *ProteinAnalysis*. The tryptophan panels in Fig. 5c, h, m were restricted to antibodies carrying exactly one arginine (R count = 1) to isolate hydrophobic contributions in the presence of minimal cationic charge. Features were computed independently for CDR H3 and VH sequences; the full feature list is provided in Section C4.

#### C3. K-mer frequency encoding

K-mer frequency features were computed using unigrams (k=1), bigrams (k=2), and trigrams (k=3), yielding a combined feature vector of approximately 8,420 dimensions (20 unigrams + 400 bigrams + 8,000 trigrams). Raw k-mer counts were normalized to frequencies summing to 1 per sequence to remove sequence length bias.

The sequence used for k-mer computation is user-configurable via the --lm flag:

| **Option** | **Sequence** | **Biological rationale** |
|---|---|---|
| --lm kmer (default) | CDR H3 | CDR3 is the primary diversity region driving polyreactivity |
| --lm kmer_vh | VH | Framework context without light chain |
| --lm kmer_vhvl | VH+VL | Full paired sequence context |

Trigrams capture local three-residue compositional motifs (such as consecutive Arg-Arg or Arg-Gly-Arg patterns) that are not captured by unigrams or bigrams alone and are biologically relevant given the dominant role of CDR H3 arginine enrichment in polyreactivity (Section A). The 8,000-dimensional trigram space is sparse for datasets of fewer than 5,000 sequences; Random Forest and XGBoost handle this sparsity natively through feature subsampling at each split (max_features=0.25 for k-mer mode). K-mer features are supported by Random Forest and XGBoost classifiers; SHAP feature importance is computed automatically after training for all encoding types including k-mer, biophysical, one-hot, and PLM embeddings (see Section F1).

#### C4. Biophysical feature encoding

Twenty-three biophysical features were computed per antibody from CDR H3 and VH sequences using the Biopython ProteinAnalysis module (Biopython 1.87). CDR H3-level features comprised: sequence length, isoelectric point (pI), net charge at pH 7.4, GRAVY hydrophobicity score, aromaticity, instability index, fraction charged residues (D+E+R+K+H), fraction hydrophobic residues (A+V+I+L+M+F+W+P), and individual counts of eleven residues of biophysical interest (Arg, Lys, His, Asp, Glu, Trp, Tyr, Phe, Gly, Pro, Cys). VH-level features comprised: full VH sequence length, VH-to-CDR3 length ratio, VH net charge at pH 7.4, and VH GRAVY hydrophobicity score. Net charge at pH 7.4 was calculated using standard pKa values for all ionizable residues (Arg, Lys, His, Asp, Glu, Cys, Tyr, N-terminus, C-terminus); pI was determined as the pH at which net charge equals zero using the Henderson–Hasselbalch equation. Biophysical features were used exclusively with Random Forest and XGBoost classifiers.

#### C5. One-hot encoding

One-hot encoding represents each antibody as a fixed-length binary matrix where each position encodes residue identity using a 20-dimensional binary indicator (one per standard amino acid). Maximum sequence lengths are fitted automatically from training data and applied consistently at inference. For Random Forest, XGBoost, and CNN, the encoded sequence is user-configurable via the --lm flag (onehot: VH+VL concatenated, 4,900 features; onehot_vh: VH only, 2,620 features; onehot_cdr3: CDR H3 only, 500 features). SHAP analysis on one-hot features attributes predictions to individual position x amino acid combinations, providing residue-level importance scores. The dual-branch TransformerOneHot encoding strategy is described in Section D.

### D TransformerOneHot: deep learning architecture for high-resolution interpretability

TransformerOneHot is a dual-branch Transformer designed specifically for position-resolved model interpretation via Integrated Gradients. Unlike PLM-based architectures that compress sequences into entangled latent vectors, TransformerOneHot encodes each antibody as a binary position x amino acid matrix, enabling IG attributions to map directly onto specific amino acids at specific sequence positions. Branch 1 encodes the concatenated VH+VL sequence (270 positions x 20 amino acids = 5,400 features; 135 VH + 135 VL, fixed maxima). Branch 2 independently encodes the CDR H3 sequence (up to 25 positions x 20 amino acids = 500 features). Both branches apply sinusoidal positional encoding and independent Pre-Layer Normalization Transformer encoder stacks, then reduce to a fixed-length vector by mean pooling. CDR3-guided cross-attention integrates the two branches: the CDR H3 summary vector serves as the query and the full VH+VL token sequence provides keys and values, with padded positions masked throughout. The combined vector feeds a classification head (LayerNorm, Dropout, Linear, softmax) producing P(Pass) and P(Fail) (Fig. S3). Training used AdamW with ReduceLROnPlateau (factor 0.5, patience 5) and gradient clipping (max_norm = 1.0). In auto mode, N layers, hidden_dim, dropout, FFN, batch size, epoch ceiling, and loss function are scaled automatically from dataset size and class balance via auto_detect_config(n, pos_rate). TransformerLM and CNN architectures are shown in Figs. S1 and S2, respectively.

### E Cross-validation, threshold optimization, and learning curves

Cross-validation used a 10-fold stratified scheme in which folds were defined by CDR H3 cluster membership (80% identity threshold; see Section A) rather than by individual antibody, keeping antibodies that share a CDR H3 cluster (>80% identity to its seed) out of opposite folds; because the clustering is greedy by seed, a small fraction of >80%-identical pairs assigned to different seeds can still span folds. This CDR H3-cluster-stratified design substantially reduces the optimistic bias that arises when highly similar sequences leak across splits, a critical safeguard given the clustered diversity of synthetic yeast display libraries. Fold composition was balanced for class labels within the cluster-stratification constraint. Model performance was evaluated using five metrics: AUC (primary metric, threshold-independent), plus accuracy, F1-score, precision, and recall at the standard 0.5 decision threshold (positive class = Pass, label = 1; non-polyreactive for PSR; monomeric for SEC). A dedicated threshold calibration module evaluated each trained model across a continuous range of decision boundaries using multiple objective functions: Youden’s J statistic (J = sensitivity + specificity − 1; maximizes the sum of true positive and true negative rates), F1-score (harmonic mean of precision and recall), F2-score (weighted F-score with β=2, emphasizing Pass recall over precision), precision-at-recall and recall-at-precision targets, and user-defined cost matrices. A pooled out-of-fold threshold was computed by aggregating predictions across all 10 folds before threshold optimization, providing a stable and reproducible deployment boundary. Separately, the representative held-out validation analysis in Fig. 3a–c,e used 2,253 antibodies (1,185 Pass / 1,068 Fail) from the fixed 80/20 split of the 11,265-antibody IPI PSR trainset; its per-antibody scores are provided in Table S4. To facilitate consistent comparison across all 25 model combinations and architectures, all manuscript performance metrics (accuracy, F1-score, precision, recall) are reported at the standard decision threshold of 0.5, enabling direct cross-model comparison without threshold-specific bias. Per-fold threshold stability and full calibration diagnostics for a representative model (TransformerLM + AbLang2, PSR) are provided in Fig. S7, demonstrating consistent performance across folds (AUC = 0.947 ± 0.007; pooled Youden threshold = 0.586; sensitivity = 0.931, specificity = 0.831 at deployment threshold). To evaluate sensitivity to framework-driven information leakage, an XGBoost classifier on AbLang2 embeddings was evaluated with entire VH germlines held out at training (leave-one-VH-germline-out, pooled out-of-germline AUC 0.903 versus 0.965 for that same classifier under CDR H3-cluster cross-validation; a dedicated re-run, cf. 0.959 in Table S1) and with VH+VL germline pairs held out (pooled AUC 0.948) (Fig. S14a). Learning curves were generated by training the TransformerLM classifier on AbLang2 embeddings over subsamples of increasing size, log-spaced from 100 antibodies to the full training set. Each size was repeated 12 times through 26,000 antibodies; DS1 sizes from 42,000 to 180,000 used four repeats, and the full 246,291-antibody requested subsample used three completed repeats; every repeat drew a fresh class-balanced subsample and a fresh CDR H3-cluster-stratified 80/20 split. Fig. 3f reports the mean and 95% CI across repeats.

### F ML Interpretability

DELPHI provides two complementary interpretability methods matched to model architecture and input representation: SHAP (SHapley Additive exPlanations) for tree-based models (Random Forest, XGBoost), and Integrated Gradients (IG) for the TransformerOneHot architecture.

SHAP TreeExplainer computes exact Shapley values by traversing the decision tree structure directly, assigning each feature a contribution that satisfies the fairness axioms of game theory (efficiency, symmetry, dummy, additivity). This is computationally exact and efficient for tree ensembles but is not applicable to neural networks, where the non-linear interaction graph is not a tree and exact Shapley computation is intractable.

We used Integrated Gradients for neural networks because it exploits the differentiability of the network: attributions are computed by backpropagating gradients from the output to the input, integrated along a straight path from a reference baseline. For the TransformerOneHot architecture, the input is a binary position x amino acid matrix, so IG attributions map directly onto specific residues at specific raw sequence positions, a biologically interpretable output. IG satisfies the completeness axiom (attributions approximate the output difference from baseline under numerical integration) and is robust to the saturation problem of simple gradient methods.

Both methods generate population-level importance profiles across all training antibodies as well as per-antibody waterfall plots, enabling interpretation at both the dataset and individual antibody level. CDR3 loop in silico mutagenesis (Section F3) provides an orthogonal perturbation analysis for feature modes that can be recomputed directly after substitution.

#### F1. SHAP interpretability (Random Forest and XGBoost)

SHAP feature importance was computed automatically after training for Random Forest and XGBoost classifiers using TreeExplainer, which provides exact SHAP values efficiently for tree-based models. SHAP analysis runs across all encoding types: k-mer frequencies, biophysical features, one-hot encoding, and PLM embeddings, with the following important distinction: when PLM embedding dimensions are present in the feature matrix, they are automatically excluded from SHAP analysis because individual PLM dimensions (emb_0…emb_N) are entangled latent features with no direct biological interpretation. SHAP is therefore computed exclusively on interpretable features (k-mer, biophysical, one-hot). If only PLM embeddings are present and no interpretable features are enabled, SHAP analysis is skipped entirely.

For the manuscript figures, SHAP values were computed on deterministic 3,000-antibody cohorts from the PSR and SEC training sets; optional subsampling bounds wall-clock time on large sets because TreeExplainer runtime scales with dataset size, number of trees and tree depth. The following outputs were generated automatically after each training run: (1) a bar chart of mean absolute SHAP values for the top 30 features, color-coded by feature type (biophysical: green; 1-mer: purple; 2-mer: violet; 3-mer: lavender; one-hot CDR H3: pink; one-hot VH/VL: orange); (2) a beeswarm plot showing the full distribution of SHAP values per feature across all samples; (3) a per-sample heatmap of SHAP values for the top 20 features; and (4) individual waterfall plots for up to 50 antibodies. A ranked CSV of mean absolute SHAP values was also exported for all features. All SHAP outputs were generated separately for training and validation sets. Regional attribution fractions were calculated by summing absolute SHAP values within each feature-defined region and normalizing within each model.

Multi-model SHAP comparison: A compare_waterfall() function enables side-by-side SHAP waterfall comparison of multiple trained models for a single antibody, displaying the cumulative SHAP contribution of the top features from each model in adjacent panels. This is used to compare different encoding strategies (for example biophysical features versus k-mer features) on the same antibody, revealing whether different representations agree on the key residue-level drivers of polyreactivity.

#### F2. Integrated Gradients (TransformerOneHot)

Integrated Gradients (IG) was used to attribute predictions of the TransformerOneHot model to individual sequence positions and amino acids. IG computes the path integral of gradients along a straight line from a reference baseline input to the actual input, providing theoretically grounded attributions that satisfy the completeness axiom in the continuous limit (the numerical approximation is checked against the prediction difference from baseline):

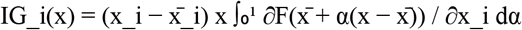

where x is the input, 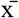 is a padding-safe, length-matched uniform amino-acid reference (1/20 in each amino-acid channel at observed positions and zero only at true padding), F is the model output P(Pass), and the integral is approximated using 200 Riemann interpolation steps. Completeness convergence was verified against 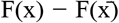; median absolute convergence deltas were 4.47 × 10^−5^ for PSR and 4.19 × 10^−5^ for SEC. Attributions were computed per antibody and accumulated as mean absolute values across all antibodies in a dataset to produce population-level importance profiles. Figure 6a,b reports mean absolute attribution at each input position; Figure 6c,d reports, for each amino acid and CDR H3 position, the mean signed attribution among antibodies carrying that residue at that position. Regional IG fractions were calculated by summing mean absolute attribution over positions within CDR H3, VH, and VL and normalizing to the total across these regions. Residue-position means were displayed only for cells represented by at least 10 antibodies.

Input structure and attribution. The TransformerOneHot dual-branch architecture encodes VH+VL sequences in branch 1 and CDR H3 independently in branch 2 (Section D). IG attributions are therefore directly interpretable: each value corresponds to a specific amino acid at a specific raw sequence position in either the VH+VL framework (branch 1) or the CDR H3 loop (branch 2). This enables simultaneous attribution to framework context and the hypervariable CDR3 region, revealing whether polyreactivity is driven primarily by CDR H3 composition, framework residues, or interactions between the two.

Outputs. The following were generated automatically: (1) population-level mean absolute IG attribution bar charts and position x amino acid heatmaps across all training antibodies, shown separately for each branch; (2) per-antibody waterfall plots for up to 50 antibodies.

#### F3. CDR3 loop in silico mutagenesis (recomputable feature modes)

CDR3 in silico mutagenesis is available for feature modes that can be recomputed directly after sequence substitution (one-hot, k-mer and biophysical encodings). For each antibody, the internal CDR H3 loop positions (sequence positions 3 through L−3) were substituted with all 20 standard amino acids while the first two and last three CDR H3 residues were held at their WT identities; each mutant was scored by the trained model. For k-mer, biophysical, and one-hot feature modes, mutant CDR3 sequences were encoded directly without re-embedding. PLM embedding mode is not supported for mutagenesis because generating a new embedding for each of the ∼20 x L mutants would require a full language model forward pass per substitution. Results were visualized as heatmaps (rows = 20 amino acids, columns = internal CDR3 loop positions labeled by their WT residues, color = ΔP(Pass) = P(Pass)mutant − P(Pass)WT on a diverging colormap centered at zero, WT residue boxed in black). This analysis identifies which CDR H3 positions and substitutions drive predicted polyreactivity risk, providing position-prioritised engineering hypotheses directly from sequence.

### G DELPHI framework and deployment

All analyses used Python 3.11.15 with NumPy 2.4.6, pandas 2.3.3, SciPy 1.17.1, scikit-learn 1.9.0, XGBoost 2.1.1, PyTorch 2.12.1, transformers 4.57.6, captum 0.9.0, SHAP 0.51.0, ablang2 0.2.1, antiberty 0.1.3, ANARCI 1.3, Biopython 1.87, Levenshtein 0.27.3 and matplotlib 3.11.0; a fixed random seed (42) was used for cross-validation splits, model training and t-SNE, and a pinned conda environment is provided in the repository. Source code, pre-trained model weights, and usage tutorials are available at https://github.com/proteininnovation/delphi.

### H Use of artificial intelligence tools

The authors independently conceived and designed DELPHI, including its configurable software framework, wrote the original Python implementation, performed the analyses, and wrote the full manuscript before using artificial intelligence tools. Large language models from OpenAI (GPT-5.4, GPT-5.5 and GPT-5.6) and Anthropic (Claude Opus 4.7, 4.8 and 5) were subsequently used to assist with code review, debugging, language editing and internal review of the manuscript. The authors evaluated and verified all AI-assisted suggestions, determined the scientific interpretation and conclusions, and take full responsibility for the accuracy and integrity of this work.

## Data availability

The public DS1 dataset is available from the original publication. The IPI PSR-ELISA and SEC training datasets are proprietary and cannot be shared publicly.

## Code availability

The DELPHI source code, documentation, public integration tests, and a worked DS1 example are available at GitHub under the MIT license. Pre-trained PSR and SEC model checkpoints are publicly available at Zenodo for immediate use with the DELPHI software. DELPHI enables users to train, evaluate, and interpret developability models using their own labelled antibody datasets. Scripts and shareable source data are provided for public reproduction of Main Figures 3, 4, and 6 and Figs. S8–S12. These workflows do not require access to the proprietary IPI training sequences.

## Acknowledgements

We thank Dr. Travis Riedel for critical reading of the manuscript. We are grateful to all members of the Institute for Protein Innovation for their support, and to the broader Institute for Protein Innovation community whose collective expertise enabled the PSR-ELISA and SEC screening campaigns that produced the datasets used in this work. This work was made possible by the generosity of Tim Springer, Chafen Lu and their family, whose gift established the endowment that supports research at the Institute for Protein Innovation.

## Author Contributions

H.N. designed and implemented the DELPHI framework, developed the code, performed and interpreted the computational analyses, provided machine-learning and technical expertise, and wrote the manuscript. A.T. conceived the project, contributed to the code and the analyses, provided technical expertise in antibody engineering, antibody development, and machine learning, and helped write the manuscript. Y.S. provided substantial revisions to the manuscript. R.C. and M.A.K. performed PSR-ELISA and SEC data curation. D.K. and H.Z. tested DELPHI PSR and SEC predictions and reviewed the manuscript. All authors read and approved the final manuscript.

