## Supplementary Information for "An interpretable open platform for sequence-based antibody developability prediction"

Contents: Figures S1–S14 and Tables S1–S6

#### Fig. S1 | TransformerLM architecture

*Fixed-length PLM embedding → 2-token Pre-LN Transformer → P(Pass)*

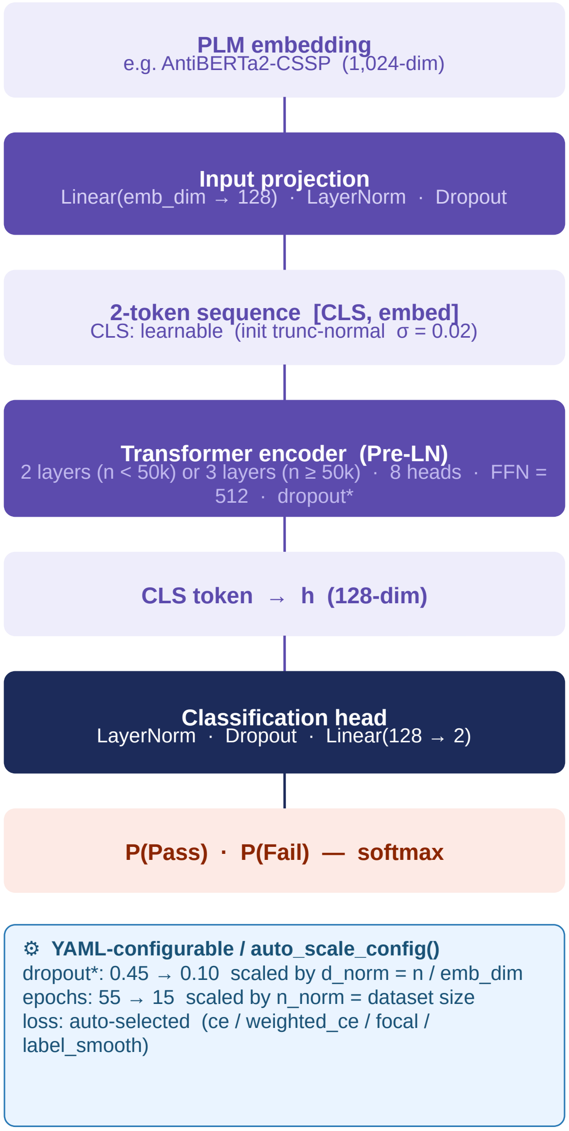


##### Fixed hyperparameters

| | |
|---|---|
| **hidden_dim** | 128 |
| **num_heads** | 8 → head_dim = 16 |
| **batch_size** | 32 |
| **lr** | $1 \times 10^{-5}$ (matched to batch = 32) |
| **FFN** | 512 (4 × hidden_dim) |
| **Pre-LN** | norm_first=True (stable at small batch) |

##### Auto-scaled parameters

| | |
|---|---|
| **dropout*** | 0.45 − 0.30 × d_norm (d_norm = samples/emb_dim) |
| **num_layers** | 2 (n < 50k) or 3 (n ≥ 50k) |
| **epochs** | 55 → 15 scaled by n_norm |
| **weight_decay** | 0.001 (n < 50k) → 0.0005 (n ≥ 50k) |
| **patience** | 7 (n < 50k) → 5 (n ≥ 50k) |

Schematic of the TransformerLM classifier. A fixed-length protein language model (PLM) embedding is projected by a Linear(emb_dim → 128) block (LayerNorm + Dropout), then combined with a learnable CLS token (truncated normal initialisation, σ = 0.02) to form a 2-token input sequence [CLS, embed]. The sequence passes through a Pre-Layer Normalisation (Pre-LN) Transformer encoder producing a CLS token representation that feeds a classification head (LayerNorm → Dropout → Linear(128 → 2)) producing P(Pass) and P(Fail) via softmax. Dashed line: the 128-dimensional CLS hidden state as a task-specific embedding for downstream clustering or t-SNE. *Dropout (0.10–0.45) and epoch ceiling (15–55) are scaled automatically by auto_scale_config() using two axes: n_norm (log-scaled dataset size) and d_norm (samples per embedding dimension). All other hyperparameters are fixed across all PLMs and dataset sizes.

#### Fig. S2 | CNN architecture

*Flat PLM embedding treated as 1D signal · dilated residual blocks · AdaptiveMaxPool*

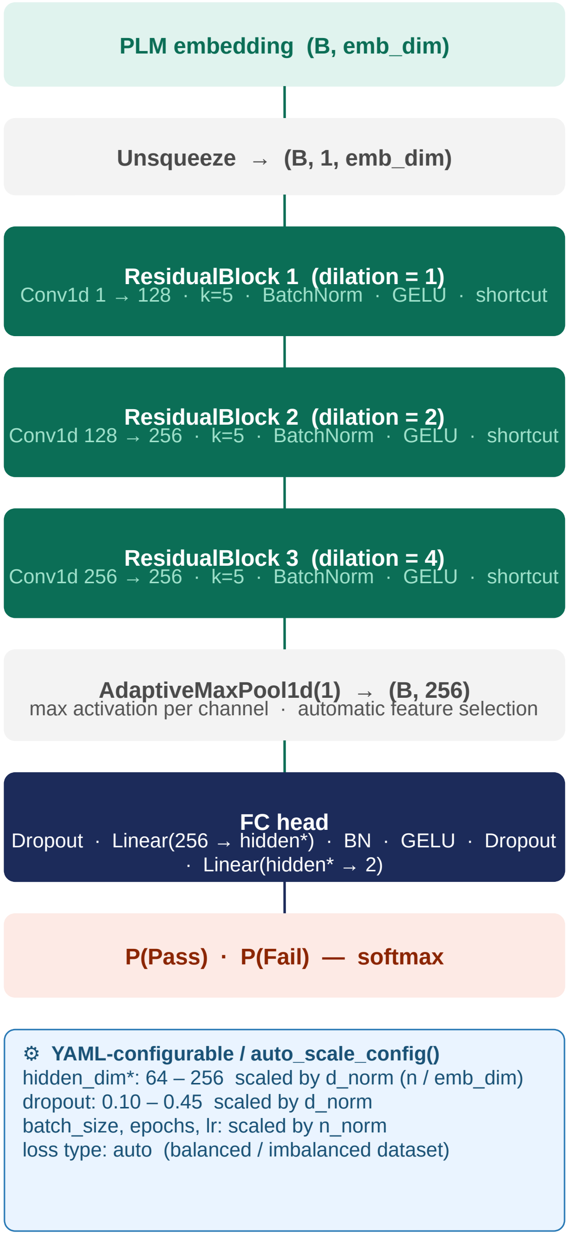


##### Architecture details

| | |
|---|---|
| **kernel_size** | 5 (all Conv1d layers) |
| **dilation** | 1, 2, 4 → cumulative receptive fields 9, 25, 57 |
| **channels** | 1 → 128 → 256 → 256 |
| **shortcut** | projection (≠ dims) or identity |
| **pooling** | AdaptiveMaxPool1d(1) → 256-dim |
| **BatchNorm** | inside every residual block |
| **activation** | GELU throughout |
| **†drop_last** | True (prevents BN crash at batch_size=1) |

##### Standalone CNN evaluation

The CNN processes the fixed-length PLM embedding vector through three dilated residual blocks, adaptive maximum pooling, and a fully connected classification head. It is evaluated as a standalone classifier in the 25-combination benchmark.

Schematic of the 1D residual CNN classifier. A PLM embedding vector (B × emb_dim) is unsqueezed to a (B, 1, emb_dim) signal and processed by three sequential dilated residual blocks (dilations 1, 2, and 4; kernel_size = 5). Each block contains two Conv1d layers, giving cumulative receptive fields of 9, 25, and 57 embedding dimensions. Each block also applies BatchNorm1d and GELU activation, with a projection shortcut where channel dimensions change. AdaptiveMaxPool1d(1) collapses the feature map to a fixed 256-dimensional vector by selecting the maximum activation per channel. The FC head applies Dropout → Linear(256 → hidden*) → BatchNorm1d → GELU → Dropout → Linear(hidden* → 2). †drop_last=True prevents BatchNorm from receiving a single-sample batch. Dashed line: penultimate-layer hidden state. *hidden_dim (64–256) and dropout (0.10–0.45) are scaled automatically by auto_scale_config() from d_norm (samples per embedding dimension).

**Fig. S3 | TransformerOneHot: Architecture and Integrated Gradients**

*Dual-branch one-hot Transformer · CDR3-guided cross-attention · position-resolved IG attribution*

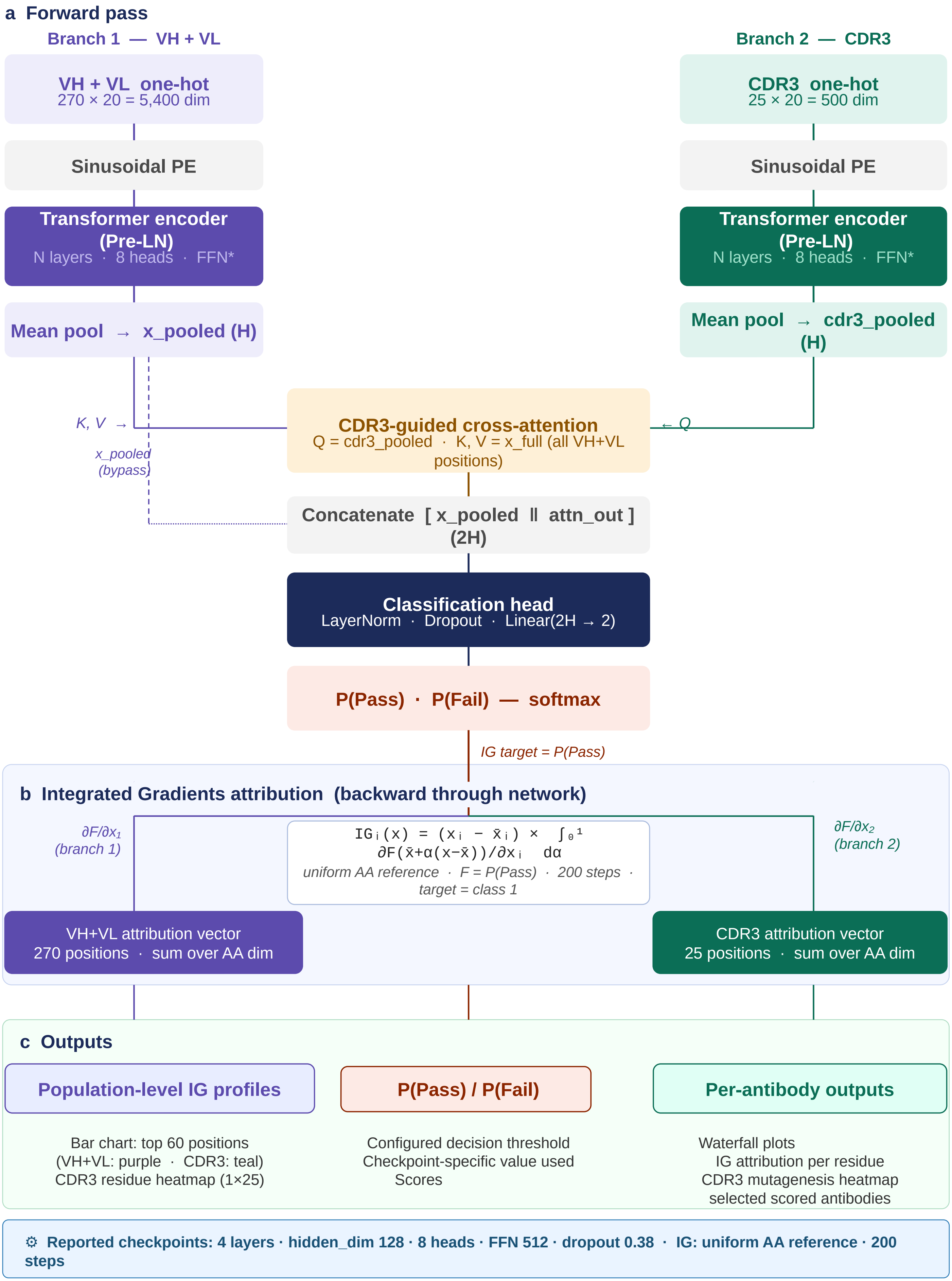




*Legend on next page*

**(a) Forward pass.** Branch 1 encodes the concatenated VH + VL sequence as a one-hot position × amino acid matrix (270 × 20 = 5,400 dimensions). Branch 2 independently encodes CDR3 (25 × 20 = 500 dimensions). Both branches apply sinusoidal positional encoding and independent Pre-LN Transformer encoder stacks, then mean-pool to x_pooled and cdr3_pooled respectively. CDR3-guided cross-attention integrates the branches: cdr3_pooled provides the query (Q) and the full VH+VL sequence provides keys and values (K, V). An x_pooled bypass (dotted line) feeds x_pooled directly to the concatenation step in parallel with the cross-attention output. The concatenated (2H)-dimensional vector feeds a classification head (LayerNorm → Dropout → Linear(2H → 2)) producing P(Pass) and P(Fail) via softmax.

**(b) Integrated Gradients attribution.** Integrated Gradients (IG) computes the path integral of gradients from a padding-safe, length-matched uniform amino-acid reference to the actual input along a straight line, approximated with 200 integration steps; attribution target = class 1 (P(Pass)). The reference assigns 1/20 to every amino-acid channel at observed positions and zero only at true padding positions. The coral arrow entering section b at the centre denotes P(Pass) as the IG attribution target. Gradients branch left ($\partial F/\partial x_1$, purple) to Branch 1 and right ($\partial F/\partial x_2$, teal) to Branch 2, propagating backward (upward arrows) through their respective encoder stacks. Summing attributions over the amino acid dimension yields a VH+VL position importance vector (270 raw sequence positions, purple box) and an CDR3 position importance vector (25 raw sequence positions, teal box).

**(c) Outputs.** Classification: P(Pass) scores thresholded at the configured, checkpoint-specific decision threshold. Population-level IG profiles (fed by VH+VL attribution vector): ranked bar chart of the top 60 positions with VH+VL positions coloured purple and CDR3 positions teal, and a 1 × CDR3_length CDR3 residue importance heatmap accumulated as mean absolute IG attributions across all antibodies. Per-antibody outputs (fed by CDR3 attribution vector): individual IG waterfall plots and CDR3 in silico mutagenesis heatmaps.

***The reported PSR and SEC checkpoints use four layers, hidden_dim 128, eight attention heads, FFN 512, dropout 0.38, and a maximum CDR3 length of 25.** SGKF, stratified group k-fold; PE, positional encoding; Pre-LN, pre-layer normalisation; IG, Integrated Gradients.

### Fig. S4 | PSR-ELISA label quality control

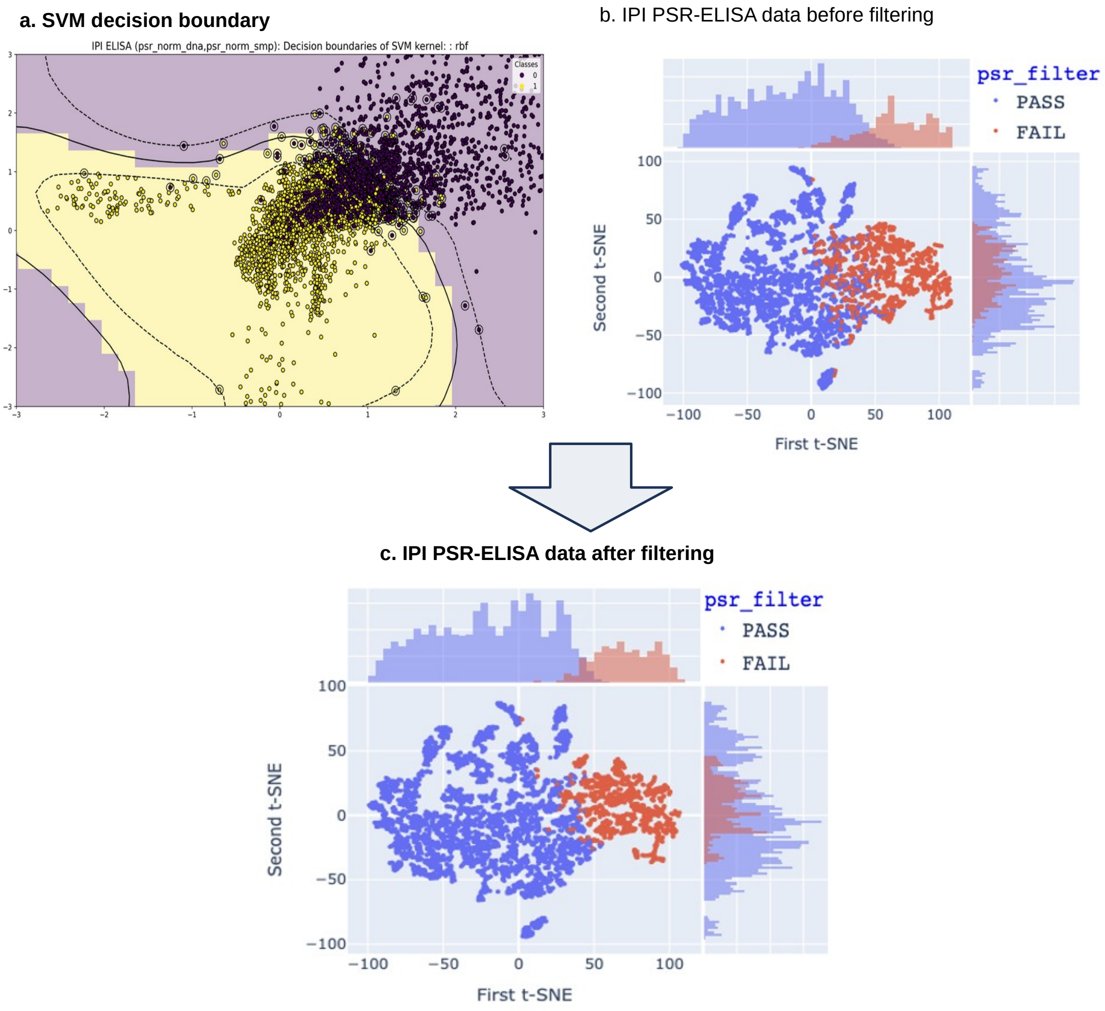


**Fig. S4 | PSR-ELISA label quality control: SVM decision-boundary modelling and t-SNE visualization.** (a) SVM decision boundary fitted to the raw DNA and OVA/SMP PSR-ELISA scores from 7,837 antibodies before quality control, with points coloured by the internally annotated Pass and Fail labels. Antibodies flagged because their Random Forest-predicted class disagreed with the annotation were concentrated near the overlapping transition region. (b) t-SNE visualization of all four continuous PSR-ELISA scores (DNA, insulin, avidin and OVA/SMP) before filtering. (c) The same t-SNE view after removal of 343 low-confidence antibodies, yielding the 7,494-antibody high-confidence set. t-SNE used perplexity = 30 and 1,000 iterations. PSR, polyspecificity reagent; SVM, support vector machine; t-SNE, t-distributed stochastic neighbour embedding; OVA, ovalbumin; SMP, soluble membrane protein.

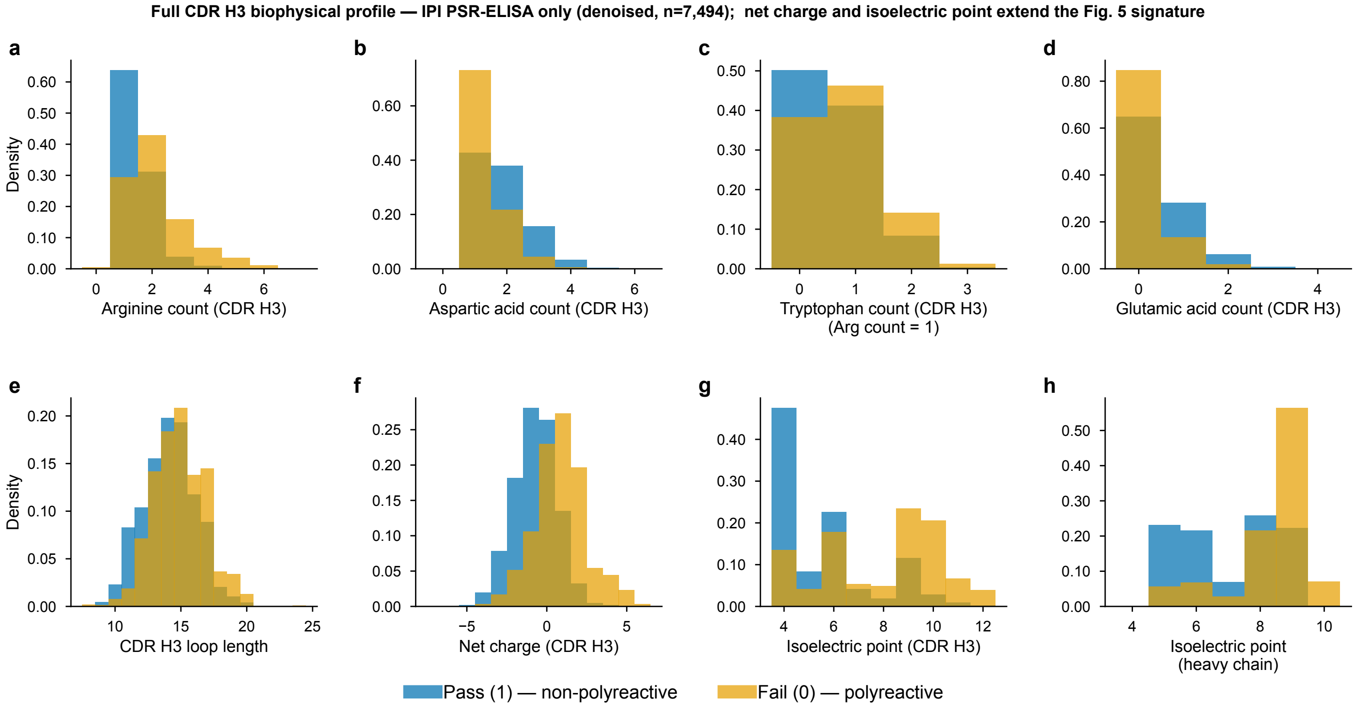


**Fig. S5 | Biophysical properties associated with polyreactivity in the IPI PSR-ELISA dataset.** Distributions in CDR H3 of (a) arginine count, (b) aspartic acid count, (c) tryptophan count (arginine count=1 subset), (d) glutamic acid count, (e) loop length, (f) net charge, (g) isoelectric point, and (h) isoelectric point of the full heavy chain variable region, for Pass (blue; low polyreactivity) and Fail (orange; high polyreactivity) antibodies (n=7,494; the ELISA-labeled subset, versus the full 11,265-antibody training set in Fig. 5). Fail antibodies show arginine enrichment, aspartic acid and glutamic acid depletion, longer loops, elevated net positive CDR H3 charge, elevated CDR H3 isoelectric point, and higher heavy-chain pI. Together, these findings indicate an association with polyreactivity. The pI measures in (g, h) are compositionally related to charge and should be interpreted as descriptive sequence properties rather than independent evidence of surface charge or a framework mechanism. Panel c is restricted to antibodies with an arginine count of 1 and should be interpreted only within this subset.

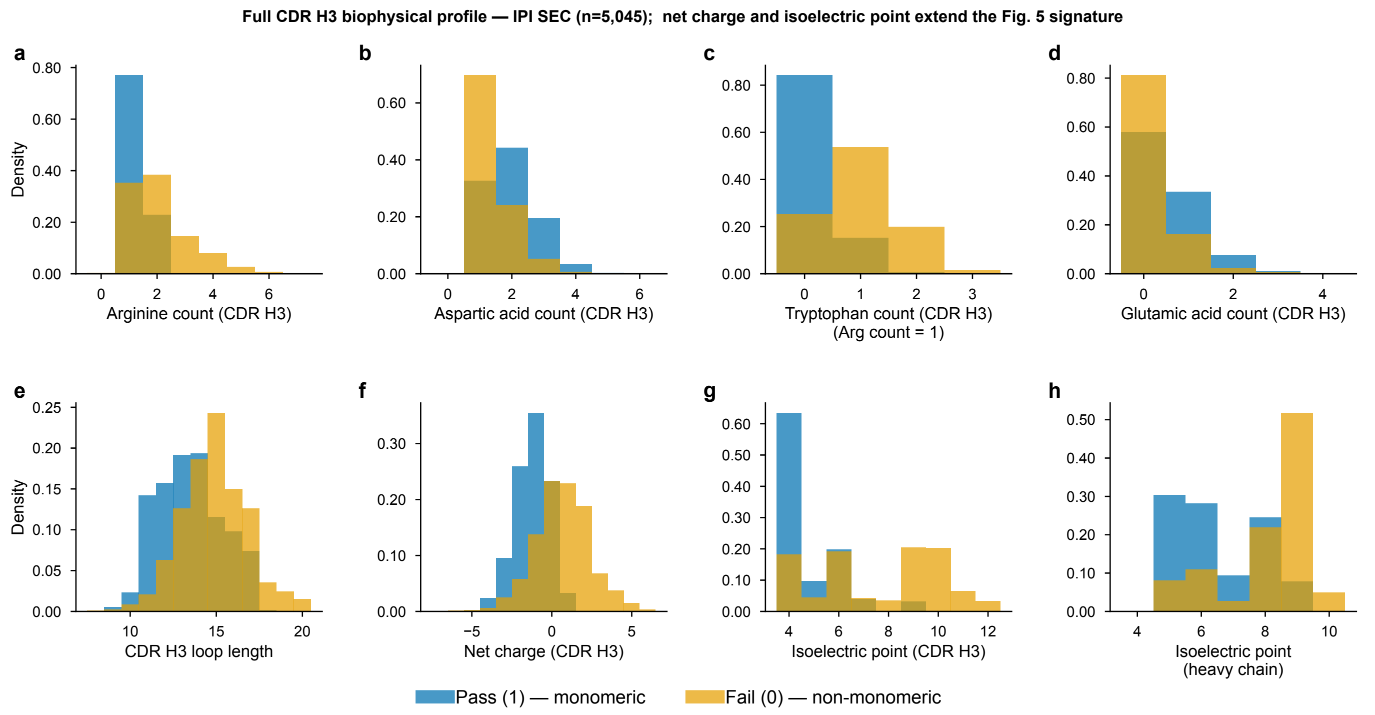


**Fig. S6 | Biophysical properties associated with SEC monomer purity failure in the IPI SEC trainset.** Distributions in CDR H3 of (a) arginine count, (b) aspartic acid count, (c) tryptophan count (arginine count=1 subset), (d) glutamic acid count, (e) loop length, (f) net charge, (g) isoelectric point, and (h) isoelectric point of the full heavy chain variable region, for Pass (blue; monomeric) and Fail (orange; non-monomeric, including aggregated or fragmented species) antibodies in the IPI SEC dataset (n = 5,045). SEC-Fail antibodies show the same directional shifts as PSR-Fail (Fig. 5a–e; Fig. S5): arginine enrichment (a), aspartate depletion (b), glutamic acid depletion (d), extended loop length (e), net positive CDR H3 charge (f), elevated CDR H3 isoelectric point (g), and higher whole-chain isoelectric point (~7.5–9 vs ~5 in Pass; h), supporting a recurring sequence association shared by polyreactivity and SEC monomer purity failure. These comparisons are descriptive within the curated SEC cohort. Because the Pass class was downsampled partly using sequence-model confidence (Methods B), effect magnitudes should be interpreted within this cohort. The pI measures in (g, h) are compositionally related to charge, and panel c is restricted to antibodies with an arginine count of 1.

**a**

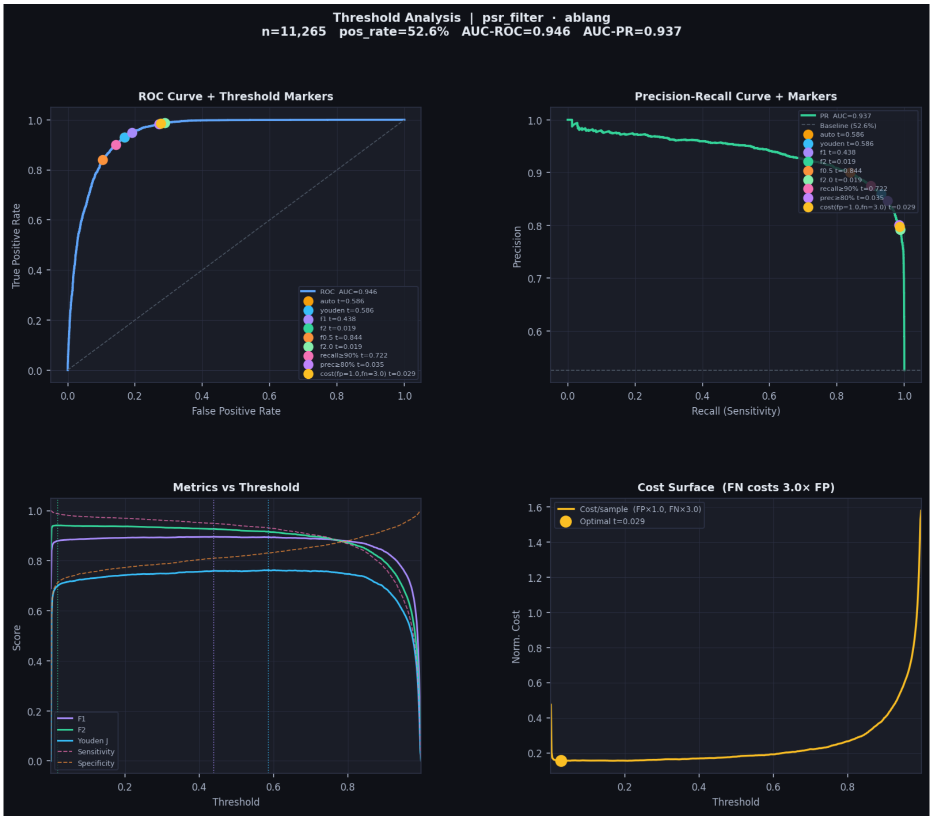


**b**

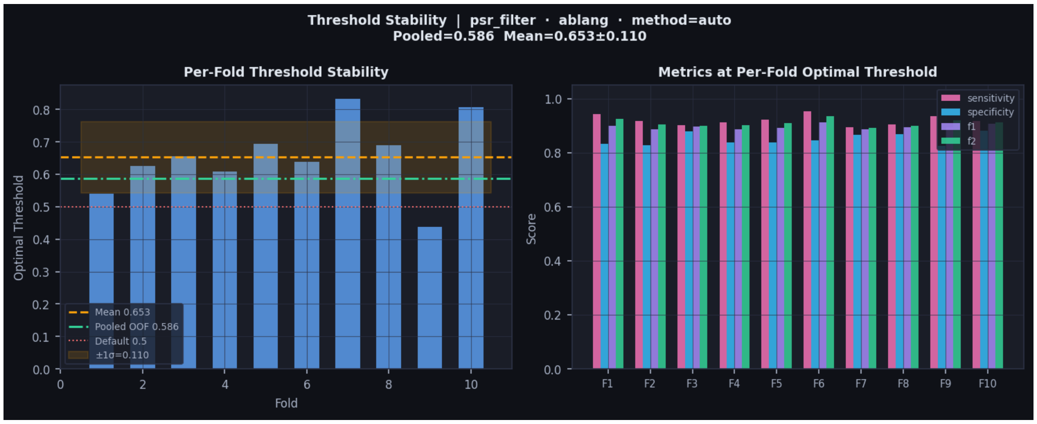


**Fig. S7 | DELPHI threshold optimization and cross-validation stability for the TransformerLM + AbLang2 PSR model (n = 11,265; 10-fold CDR H3-stratified cross-validation).** (a) Four-panel threshold analysis on pooled out-of-fold predictions. Top left: ROC curve (AUC-ROC = 0.946) with markers for seven threshold-selection criteria: Youden's J (t = 0.586; also selected by the automatic rule), F1 (t = 0.438), F2 (t = 0.019), F0.5 (t = 0.844), recall at least 90% (t = 0.722), precision at least 80% (t = 0.035), and a cost-sensitive criterion with FP = 1 and FN = 3 (t = 0.029). Top right: precision-recall curve (AUC-PR = 0.937) with the same markers. Bottom left: F1, F2, Youden's J, sensitivity and specificity across thresholds. Bottom right: normalized cost surface for the stated cost ratio. (b) Per-fold threshold stability across 10 folds. Left: fold-specific Youden thresholds (bars), their mean (orange dashed; t = 0.653 ± 0.110), pooled out-of-fold threshold (green dash-dot; t = 0.586), and reference threshold 0.5 (red dotted). Right: sensitivity, specificity, F1 and F2 at each fold-specific threshold. At the pooled threshold, sensitivity = 0.931, specificity = 0.831, F1 = 0.894 and AUC-ROC = 0.946. PR, precision-recall; OOF, out-of-fold; FP, false positive; FN, false negative.

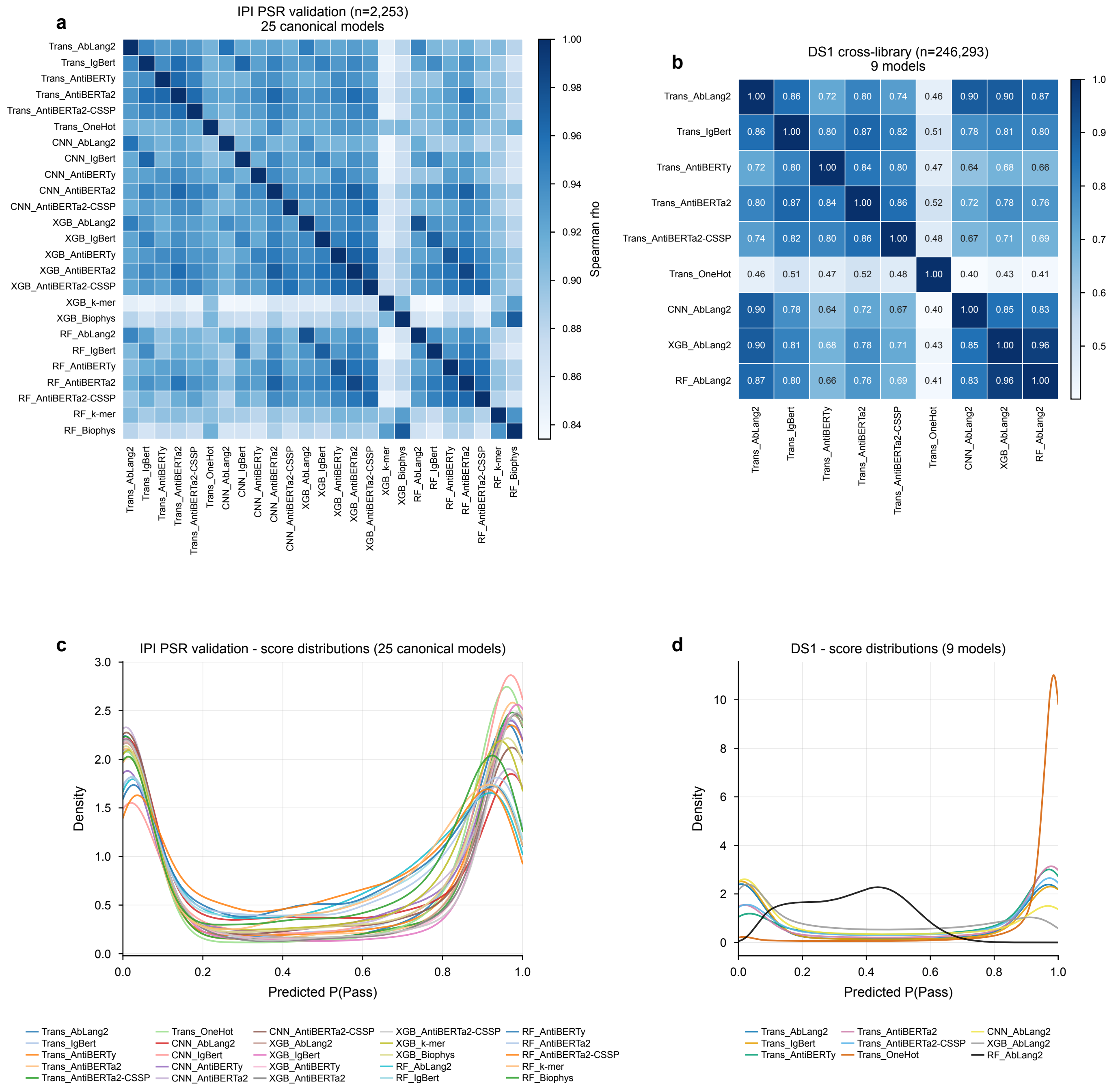


**Fig. S8 | Cross-model prediction-score correlations and score-distribution shapes on within-distribution and cross-library evaluation sets.** All 25 model combinations were trained on the IPI PSR trainset (80%, n = 9,012) and evaluated in two settings: the held-out IPI PSR validation split (20%, n = 2,253; panels a, c) and the full DS1 public dataset (n = 246,293; cross-library transfer; panels b, d). (a) Spearman's ρ between predicted P(Pass) scores for all 25 models on IPI validation. PLM-based models cluster tightly (within-architecture ρ = 0.91–0.97; between-architecture ρ = 0.88–0.96), while count- and physicochemical-feature baselines (k-mer, biophysical) form a distinct low-correlation block (ρ ≈ 0.83–0.90). (b) Spearman's ρ between nine models on DS1: six Transformers plus CNN, XGBoost, and RF paired with AbLang2. The three non-Transformer AbLang2 models stay highly correlated with Transformer+AbLang2 (ρ = 0.87–0.91), showing similar score rankings across classifier architectures that share AbLang2 embeddings. Transformer+OneHot is the sole outlier (ρ = 0.40–0.52 with every PLM-based model); one-hot encoding produces a distinct cross-library score ranking from the learned PLM representations. (c) KDE of predicted P(Pass) on IPI PSR validation set (25 models). All PLM-based models produce clean bimodal distributions, showing within-distribution score separation and agreement among models. (d) KDE of predicted P(Pass) on full DS1 (9 models). Transformer+OneHot (brown) collapses with peak density above 11 at P(Pass) ≈ 1, the signature of score collapse toward the majority class under distribution shift, providing a visual correlate of the quantitative transfer gap in Fig. 4a and Table S2. Non-Transformer AbLang2 models show broader, less bimodal shapes consistent with the cross-library accuracy drop in Table S2.
Spearman's ρ computed on raw scores; KDE bandwidth = 0.15.

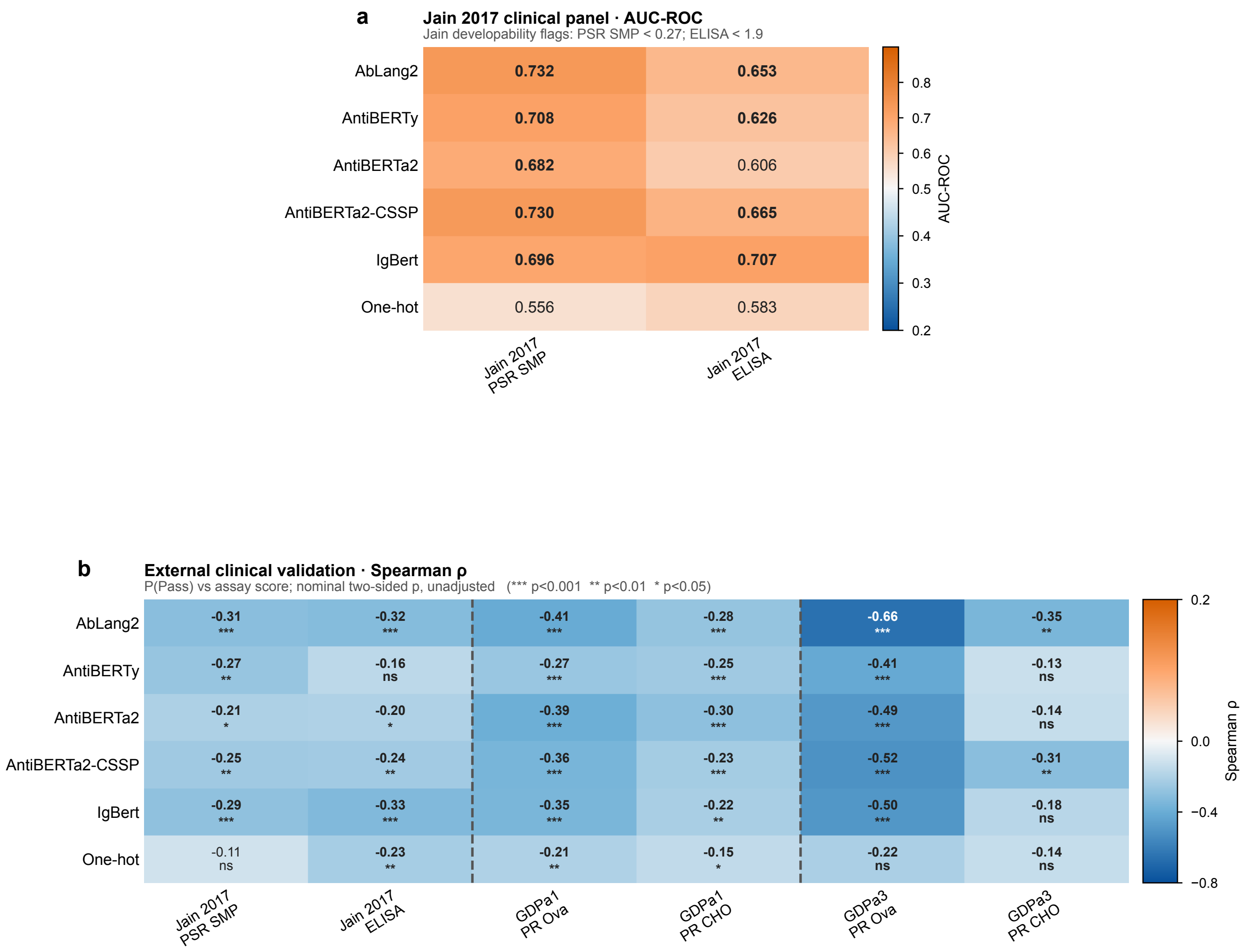


**Fig. S9 | Full external clinical-validation grid.** Zero-shot evaluation of IPI-PSR-trained models on three external clinical-stage cohorts. (a) AUC-ROC for six language models (AbLang2, AntiBERTy, AntiBERTa2, AntiBERTa2-CSSP, IgBert, one-hot) on the two Jain 2017 clinical-panel readouts (PSR-SMP and ELISA), each binarized by its own Jain developability flag (PSR score < 0.27; ELISA < 1.9); AbLang2 shows comparatively strong point estimates across these readouts. The Ginkgo cohorts (GDPa1, GDPa3) report PR-CHO/PR-Ova on a different, bead-based scale and are evaluated by rank correlation only (panel b), not a binary threshold. (b) Spearman ρ between predicted P(Pass) and the continuous assay score, with significance (*** $p < 0.001$, ** $p < 0.01$, * $p < 0.05$, ns otherwise); correlations are negative as expected for P(Pass) versus polyreactivity, strongest for AbLang2 on GDPa3 PR-Ova ($\rho = -0.66$, on a small $n = 80$ cohort). Main Fig. 4b–d and f focus on the deployed Transformer + AbLang2 model, whereas Main Fig. 4e compares the five PLM-based Transformer models on GDPa3 PR-CHO. Panel values are point estimates without confidence intervals and are intended for descriptive comparison rather than fine model ranking. Reported P values are nominal, two-sided, and unadjusted for multiple comparisons.

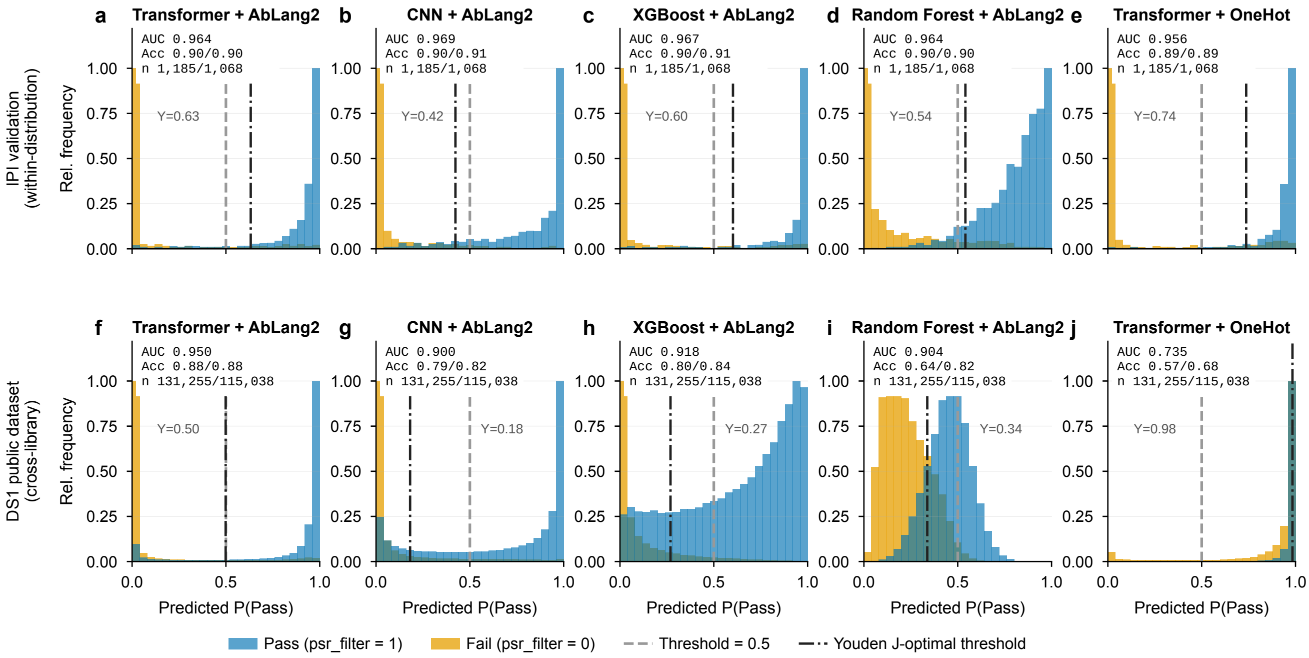


**Fig. S10 | Predicted-score distributions per architecture, within-distribution versus cross-library.** Predicted P(Pass) histograms for Pass (blue) and Fail (orange) antibodies for five architectures, Transformer + AbLang2, CNN + AbLang2, XGBoost + AbLang2, Random Forest + AbLang2, and Transformer + one-hot, on the held-out IPI PSR validation split (top row, a–e; n = 2,253) and on the full DS1 public library (bottom row, f–j; n = 246,293). Each panel annotates AUC, accuracy at the 0.5 and Youden-J thresholds, and class sizes; the dashed line marks the 0.5 threshold and the dash-dot line the Youden-J threshold. Within distribution all five displayed architectures are sharply bimodal (score separation, not formal calibration, which is shown in Fig. 3c); on DS1 the one-hot Transformer collapses toward P(Pass) ≈ 1, the visual correlate of the cross-library transfer gap (Fig. 4a; Table S2).

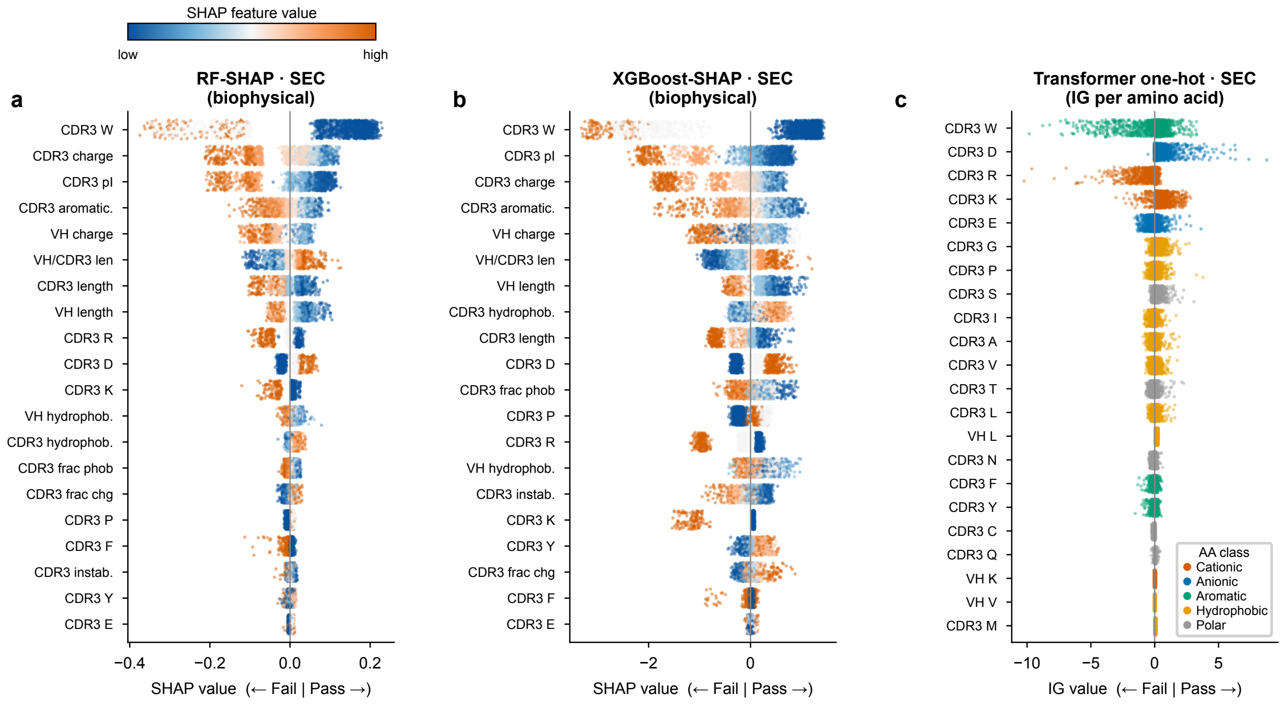


**Fig. S11 | Cross-model interpretability of SEC monomer purity failure predictions.** (a, b) SHAP beeswarm plots for Random Forest (a) and XGBoost (b) trained on biophysical CDR3 and VH features. Each dot represents one antibody; x-position = signed SHAP value (negative = Fail-associated attribution, positive = Pass-associated attribution); dot colour = feature value (blue = low, red = high). Features are ordered by mean |SHAP|. (c) Integrated Gradients (IG) attributions from the one-hot Transformer, showing the top 22 (region, amino-acid) rows ranked by mean |IG|, with CDR H3 and VH framework residues interleaved by importance (the row-label prefix gives the region). Dot colour indicates AA physicochemical class, as in the panel legend. The two biophysical models (a, b) rank CDR3 tryptophan content as the top feature for SEC, ahead of net charge, isoelectric point, and aromaticity, while the one-hot Transformer (c) places tryptophan among the leading risk-driving residues alongside arginine (R), with aspartate (D) protective. The Arg-risk/Asp-protective attribution pattern is directionally consistent with the PSR analysis (Fig. S12), while tryptophan carries the largest single-feature SHAP weight for SEC. Because correlated descriptors divide attribution and the SEC Pass class was selected partly using sequence-model confidence (Methods B), feature ranks are model-specific. Agreement across these models trained on the same selected dataset is computational recurrence, not independent biological validation. SHAP panels use a deterministic n = 3,000 sample from the IPI SEC trainset; Transformer IG uses all n = 5,045 labeled antibodies (target = Pass, 200 integration steps, length-matched uniform amino-acid reference). Panels show up to 1,500 points per feature row for rendering clarity; SHAP feature ranking uses the deterministic 3,000-antibody SHAP cohort, whereas Transformer IG ranking uses all labeled antibodies. SHAP, SHapley Additive exPlanations; IG, Integrated Gradients; CDR H3, heavy chain complementarity-determining region 3; pI, isoelectric point; CDR3, complementarity-determining region 3; AA, amino acid.

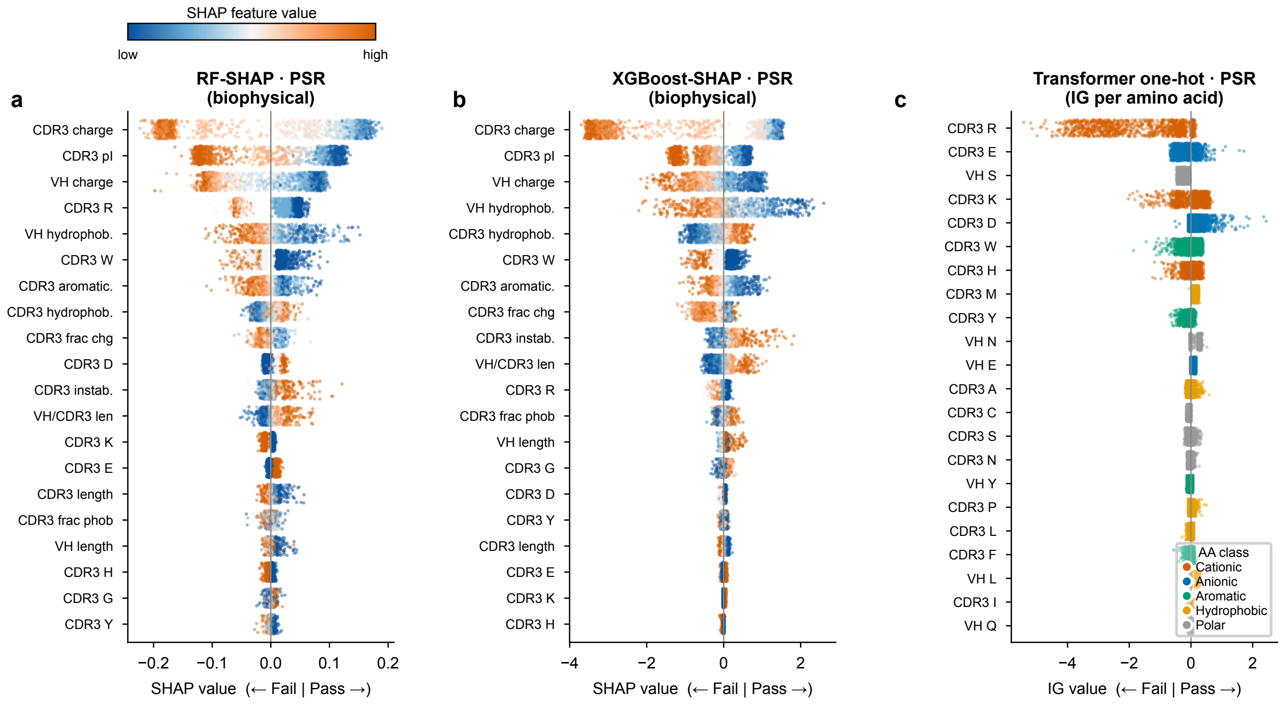


**Fig. S12 | Cross-model interpretability of polyreactivity (PSR-filter) predictions.** (a, b) SHAP beeswarm plots for Random Forest (a) and XGBoost (b) models trained on biophysical features derived from CDR H3 and VH sequences. Each dot represents one antibody; x-position indicates the signed SHAP value (negative = pushes prediction toward FAIL; positive = toward PASS); dot colour encodes the feature value (blue = low, red = high). Features are ranked by mean |SHAP| (top = most important). Panels a and b are ordered independently, each by its own mean |SHAP|. (c) Per-amino-acid Integrated Gradients (IG) attributions from the one-hot sequence Transformer, showing the top 22 (region, amino-acid) rows ranked by mean |IG|, with CDR H3 and VH framework residues interleaved by importance (the row-label prefix gives the region). Each row is one amino acid in one region, and each dot is one antibody's IG summed over that residue's positions in that region. Dot colour indicates AA physicochemical class (red = cationic R,K,H; blue = anionic D,E; green = aromatic W,F,Y; orange = hydrophobic A,G,I,L,M,P,V; grey = polar S,T,N,Q,C). Negative IG = Fail-associated attribution; positive IG = Pass-associated attribution. Across the three models, aggregate-feature SHAP and residue-level IG show directionally compatible attribution patterns for cationic and anionic composition and tryptophan. Because charge, pI, and residue-count features are correlated and mathematically related, SHAP feature ranks are model-specific and should not be read as a unique hierarchy of biological mechanisms. These analyses provide computational consistency across models trained on the same labels, not independent experimental validation. SHAP panels use a deterministic n = 3,000 sample from the IPI PSR trainset; Transformer IG uses all n = 11,265 labeled antibodies (target = Pass, 200 integration steps, length-matched uniform amino-acid reference). Panels show up to 1,500 points per feature row for rendering clarity; SHAP feature ranking uses the deterministic 3,000-antibody SHAP cohort, whereas Transformer IG ranking uses all labeled antibodies. SHAP, SHapley Additive exPlanations; IG, Integrated Gradients; CDR H3, heavy chain complementarity-determining region 3; pI, isoelectric point; CDR3, complementarity-determining region 3; AA, amino acid.

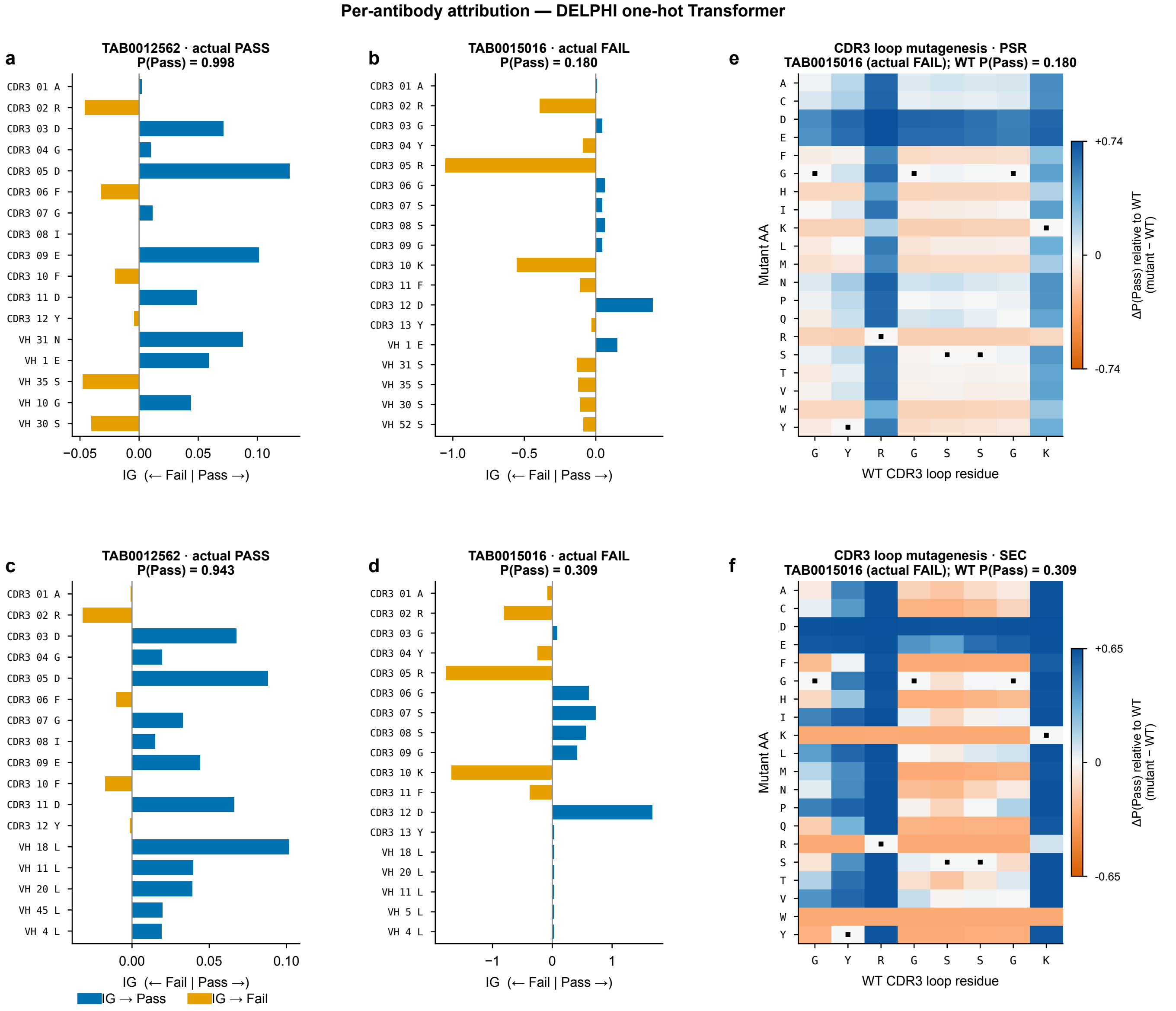


**Fig. S13 | Residue-level per-antibody interpretation using TransformerOneHot + Integrated Gradients.** Two IPI antibodies with experimental PSR and SEC Pass/Fail labels are shown together with their TransformerOneHot model scores: TAB0012562 is labeled Pass in both assays (PSR P(Pass) = 0.998; SEC P(Pass) = 0.943); TAB0015016 is labeled Fail in both assays (PSR P(Pass) = 0.180; SEC P(Pass) = 0.309; both below the deployed Pass decision threshold and therefore correctly classified Fail). All three DELPHI models (RF-biophysical, XGBoost-biophysical, TransformerOneHot) correctly classify both antibodies, providing illustrative examples for attribution analysis. Top row (PSR): (a) IG profile of the Pass antibody. CDR3_02_R shows moderate risk-increasing attribution (orange) but is balanced by multiple protective anionic residues (D3, D5, E9, D11; blue), showing that the model attribution reflects sequence context rather than arginine presence alone. (b) IG profile of the Fail antibody. CDR3_02_R and CDR3_05_R show strong risk-increasing attribution (orange), joined by CDR3_10_K; CDR3_12_D has a Pass-directed attribution. (e) CDR3 loop in silico mutagenesis heatmap for the PSR-Fail antibody. Each of the 20 amino acids is substituted at every CDR3 loop residue and rescored with the complete antibody sequence; colour encodes ΔP(Pass) relative to WT: blue = increased P(Pass), orange = decreased P(Pass), and white = no change; black squares mark WT residues (Δ = 0), and the x-axis reports the WT CDR3 loop residues. Anionic or neutral substitutions at the loop arginine and lysine residues consistently increase ΔP(Pass) relative to WT, providing residue-level computational engineering hypotheses. Bottom row (SEC): (c, d) IG profiles for the Pass and Fail antibodies mirror the PSR pattern; the Fail antibody shows larger absolute cationic attribution under SEC than PSR, consistent with the shared CDR H3 electrostatic liability. (f) CDR3 loop mutagenesis under the SEC model predicts that the same substitutions increase ΔP(Pass) relative to WT for both liabilities. IG used the same Pass-class target, 200 integration steps, and padding-safe length-matched uniform amino-acid reference as the cohort analysis. Per-antibody signed IG reports the contribution assigned to each observed residue in that antibody and reflects both model-learned residue-position patterns and sequence-context-dependent interactions. These panels show model attributions and in silico score perturbations; they do not experimentally establish improved expression, binding, PSR, or SEC behavior.

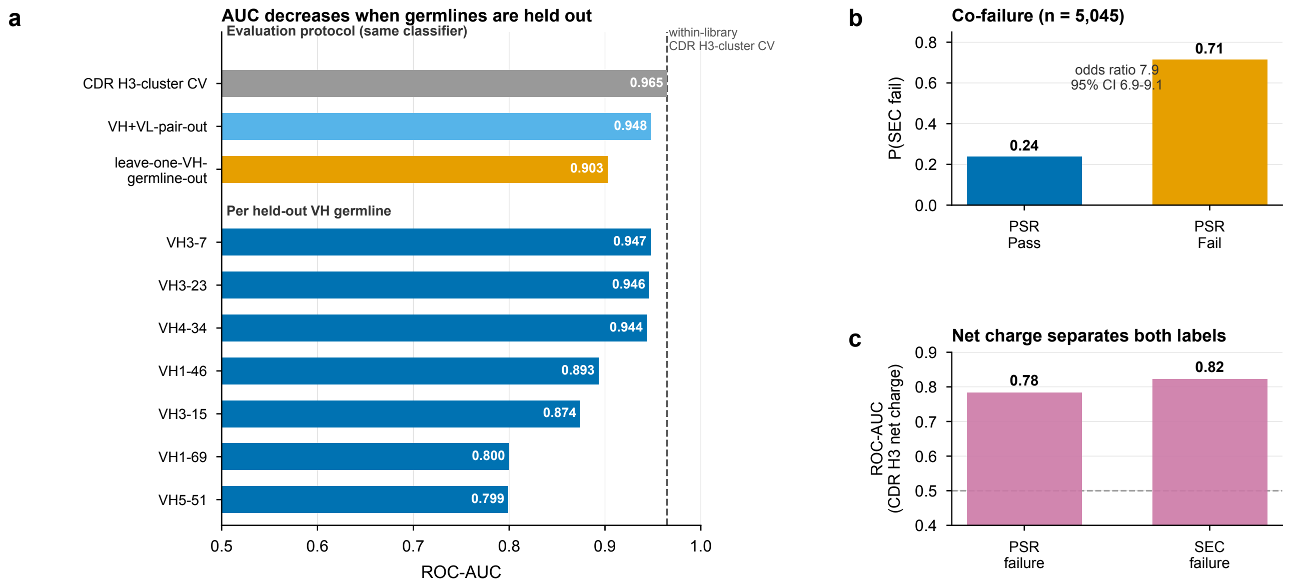


**Fig. S14 | Framework-grouped evaluation and a molecule-level cross-assay test.** (a) ROC-AUC of an XGBoost classifier on AbLang2 embeddings on the IPI PSR trainset (n = 11,265) under three evaluation protocols with the identical classifier: 10-fold CDR H3-cluster-stratified cross-validation (pooled out-of-fold), leave-one-VH-germline-out, and leave-one-VH+VL-germline-pair-out (each pooled across held-out folds); the dashed line marks the within-library CDR H3-cluster cross-validation value. Lower bars give per-held-out-VH-germline AUC for the seven germlines with $n \geq 100$. (b) Among the 5,045 antibodies with both PSR and SEC labels, the fraction failing SEC for PSR-Pass versus PSR-Fail antibodies (odds ratio and 95% confidence interval annotated; 2×2 Fisher exact $p < 10^{-200}$). (c) ROC-AUC of CDR H3 net charge alone (a computed biophysical feature, not a model score) for discriminating PSR and SEC failure labels within the same molecules; dashed line marks chance. Pass and Fail are the experimental assay calls.

**Table S1 | IPI PSR Dataset;10-Fold CDR H3-Stratified Cross-Validation; All Model Combinations.** Classification performance (accuracy, precision, recall, F1-score, AUC-ROC) for all 25 model combinations: five classifier architectures (CNN, Transformer, Random Forest [RF], XGBoost, and a dual-branch one-hot Transformer): the first four paired with five antibody-specific protein language models (AbLang2, AntiBERTy, AntiBERTa2, AntiBERTa2-CSSP, IgBert), the fifth applied to one-hot encoding, and RF and XGBoost additionally with two traditional feature baselines (biophysical descriptors; k-mer frequency encodings), under 10-fold CDR H3-cluster-stratified cross-validation on the IPI PSR trainset (n = 11,265; ~53%/47% Pass/Fail). Positive class = Pass (label = 1; low polyreactivity; non-polyreactive). Accuracy, precision, recall, and F1-score are reported at standard threshold = 0.5; AUC is threshold-independent. PLM, protein language model; RF, Random Forest; AUC, area under the receiver operating characteristic curve; CDR H3, heavy chain complementarity-determining region 3; SGKF, stratified group k-fold.

| Architecture | Language Model | Accuracy | Precision | Recall | F1-Score | AUC |
|---|---|---|---|---|---|---|
| CNN | AbLang2 | 0.895 | 0.875 | 0.933 | 0.903 | 0.959 |
| | AntiBERTy | 0.896 | 0.875 | 0.936 | 0.904 | 0.963 |
| | AntiBERTa2 | 0.902 | 0.887 | 0.932 | 0.909 | 0.964 |
| | AntiBERTa2-CSSP | 0.901 | 0.889 | 0.938 | 0.908 | 0.963 |
| | IgBert | 0.904 | 0.905 | 0.904 | 0.904 | 0.964 |
| Transformer | AbLang2 | 0.883 | 0.887 | 0.883 | 0.882 | 0.946 |
| | AntiBERTy | 0.897 | 0.899 | 0.897 | 0.897 | 0.961 |
| | AntiBERTa2 | 0.902 | 0.890 | 0.929 | 0.909 | 0.964 |
| | AntiBERTa2-CSSP | 0.893 | 0.894 | 0.893 | 0.892 | 0.958 |
| | IgBert | 0.899 | 0.901 | 0.899 | 0.899 | 0.960 |
| | One-hot | 0.903 | 0.888 | 0.934 | 0.906 | 0.961 |
| RF | AbLang2 | 0.890 | 0.855 | 0.943 | 0.897 | 0.952 |
| | AntiBERTy | 0.887 | 0.858 | 0.931 | 0.893 | 0.954 |
| | AntiBERTa2 | 0.896 | 0.867 | 0.939 | 0.901 | 0.959 |
| | AntiBERTa2-CSSP | 0.882 | 0.854 | 0.942 | 0.889 | 0.948 |
| | IgBert | 0.897 | 0.873 | 0.941 | 0.906 | 0.961 |
| | Biophysical | 0.881 | 0.858 | 0.927 | 0.891 | 0.945 |
| | k-mer | 0.876 | 0.851 | 0.927 | 0.888 | 0.939 |
| XGBoost | AbLang2 | 0.897 | 0.875 | 0.931 | 0.902 | 0.959 |
| | AntiBERTy | 0.896 | 0.879 | 0.924 | 0.901 | 0.961 |
| | AntiBERTa2 | 0.903 | 0.884 | 0.931 | 0.907 | 0.963 |
| | AntiBERTa2-CSSP | 0.898 | 0.881 | 0.925 | 0.902 | 0.961 |
| | IgBert | 0.903 | 0.891 | 0.929 | 0.909 | 0.967 |
| | Biophysical | 0.876 | 0.859 | 0.915 | 0.886 | 0.946 |
| | k-mer | 0.879 | 0.873 | 0.917 | 0.888 | 0.942 |

**Table S2** | **Cross-Dataset Generalization of Transformer Architecture**

Five-metric classification performance (accuracy, precision, recall, F1-score, AUC-ROC) for the Transformer architecture paired with six sequence encodings (ABLang, AntiBERTy, AntiBERTa2, AntiBERTa2-CSSP, IgBERT, one-hot) under three evaluation regimes: (i) DS1 10-fold CDR H3-cluster-stratified cross-validation (internal; train and test on DS1, n = 246,293); (ii) DS1→IPI transfer (train on DS1, evaluate on IPI PSR trainset, n = 11,265); and (iii) IPI→DS1 transfer (train on IPI PSR trainset, evaluate on DS1, n = 246,293). Positive class = Pass (label = 1; low polyreactivity/non-polyreactive). Accuracy, precision, recall, and F1-score are reported at standard threshold = 0.5; AUC is threshold-independent. DS1, public VH-domain antibody dataset (1); IPI, Institute for Protein Innovation; PLM, protein language model; AUC, area under the receiver operating characteristic curve; CDR H3, heavy-chain complementarity-determining region 3.

| Architecture | Language Model | Accuracy | Precision | Recall | F1-Score | AUC |
|---|---|---|---|---|---|---|
| **DS1 · 10-Fold Cross-Validation (Train DS1, Test DS1)** | | | | | | |
| Transformer | ABLang | **0.9478** | 0.9484 | **0.9478** | **0.9478** | 0.9850 |
| Transformer | AntiBERTa2-CSSP | 0.9348 | 0.9358 | 0.9348 | 0.9347 | 0.9806 |
| Transformer | One-hot | 0.9366 | **0.9547** | 0.9278 | 0.9390 | **0.9882** |
| **DS1 → IPI Transfer (Train DS1, Predict IPI)** | | | | | | |
| Transformer | ABLang | **0.7862** | **0.7765** | 0.8336 | **0.8040** | 0.8526 |
| Transformer | AntiBERTy | 0.7066 | 0.7430 | 0.6761 | 0.7080 | 0.7685 |
| Transformer | AntiBERTa2 | 0.7406 | 0.7464 | 0.7676 | 0.7569 | 0.7828 |
| Transformer | AntiBERTa2-CSSP | 0.6762 | 0.6872 | 0.7055 | 0.6962 | 0.7407 |
| Transformer | IgBERT | 0.7223 | 0.6567 | **0.9894** | 0.7894 | **0.8833** |
| Transformer | One-hot | 0.7119 | 0.7663 | 0.6508 | 0.7038 | 0.7886 |
| **IPI → DS1 Transfer (Train IPI, Predict DS1)** | | | | | | |
| Transformer | ABLang | **0.8806** | **0.9012** | 0.8715 | **0.8861** | **0.9501** |
| Transformer | AntiBERTy | 0.7928 | 0.7466 | 0.9252 | 0.8264 | 0.8923 |
| Transformer | AntiBERTa2 | 0.8319 | 0.7825 | 0.9481 | 0.8574 | 0.9313 |
| Transformer | AntiBERTa2-CSSP | 0.8182 | 0.7900 | 0.8973 | 0.8402 | 0.9038 |
| Transformer | IgBERT | 0.8588 | 0.8965 | 0.8309 | 0.8625 | 0.9369 |
| Transformer | One-hot | 0.5720 | 0.5550 | **0.9935** | 0.7122 | 0.7351 |

**Table S3 | SEC monomer purity failure prediction performance on the IPI SEC training set**. Classification performance of DELPHI models for SEC monomer purity failure prediction (5,045 antibodies; ~64% Pass [monomeric] / 36% Fail [non-monomeric]). Overall accuracy, precision, recall and F1-score are reported under 10-fold CDR H3-cluster-stratified cross-validation for 20 architecture–PLM combinations spanning Transformer, CNN, XGBoost and Random Forest paired with AntiBERTa2-CSSP, AntiBERTa2, AntiBERTy, AbLang2 and IgBert, plus Transformer One-Hot encoding, and IPI-specific models using Biophysical CDR3 features and k-mer sequence features with RF and XGBoost. Positive class = Pass (label = 1; monomeric).

| Architecture | Language Model | Accuracy | Precision | Recall | F1-Score | AUC |
|---|---|---|---|---|---|---|
| CNN | AbLang2 | 0.8547 | 0.8542 | 0.9306 | 0.8906 | 0.9124 |
| | AntiBERTy | 0.901 | 0.904 | 0.901 | 0.9 | 0.952 |
| | AntiBERTa2 | 0.874 | 0.882 | 0.874 | 0.873 | 0.94 |
| | AntiBERTa2-CSSP | 0.876 | 0.883 | 0.876 | 0.876 | 0.943 |
| | IgBert | 0.808 | 0.85 | 0.808 | 0.8 | 0.93 |
| Transformer | AbLang2 | 0.8234 | 0.8018 | 0.8234 | 0.8061 | 0.8769 |
| | AntiBERTy | 0.8963 | 0.8994 | 0.8963 | 0.8939 | 0.9381 |
| | AntiBERTa2 | 0.8993 | 0.9013 | 0.8993 | 0.8973 | 0.9386 |
| | AntiBERTa2-CSSP | 0.8940 | 0.8963 | 0.8940 | 0.892 | 0.9402 |
| | IgBert | 0.863 | 0.841 | 0.863 | 0.8475 | 0.9079 |
| | One-Hot | 0.9241 | 0.9012 | 0.9897 | 0.9433 | 0.9566 |
| RF | AbLang2 | 0.8547 | 0.8542 | 0.9306 | 0.8906 | 0.9124 |
| | AntiBERTy | 0.8924 | 0.8896 | 0.9486 | 0.9181 | 0.9439 |
| | AntiBERTa2 | 0.8789 | 0.8793 | 0.9392 | 0.9080 | 0.9357 |
| | AntiBERTa2-CSSP | 0.8616 | 0.8593 | 0.9364 | 0.8959 | 0.9233 |
| | IgBert | 0.8805 | 0.8760 | 0.946 | 0.9096 | 0.9342 |
| | Biophysical | 0.9336 | 0.9228 | 0.9775 | 0.9467 | 0.9573 |
| | Kmer | 0.9388 | 0.9295 | 0.9779 | 0.9530 | 0.9581 |
| XGBoost | AbLang2 | 0.8826 | 0.8878 | 0.9327 | 0.9096 | 0.9349 |
| | AntiBERTy | 0.9169 | 0.9185 | 0.9542 | 0.9360 | 0.9561 |
| | AntiBERTa2 | 0.9090 | 0.9133 | 0.9474 | 0.9298 | 0.9524 |
| | AntiBERTa2-CSSP | 0.9048 | 0.9077 | 0.9470 | 0.9268 | 0.9468 |
| | IgBert | 0.9108 | 0.9125 | 0.9511 | 0.9313 | 0.9509 |
| | Biophysical | 0.9342 | 0.9295 | 0.9704 | 0.9494 | 0.9596 |
| | Kmer | 0.9293 | 0.9412 | 0.9646 | 0.9527 | 0.9366 |

**Table S4 | PSR prediction score of validation data (train:80, validation:20%) of IPI PSR trainset and DS1**

**Table S5 | PSR prediction scores of clinical antibodies**

**Table S6 | Comparison of DELPHI with recent sequence-based machine-learning methods for antibody developability prediction.**
Leakage-aware cross-validation denotes sequence-similarity-stratified splitting: Amini et al. use whole-sequence clusters and DELPHI uses CDR H3 clusters, whereas Abeer et al. select a diverse hold-out test set by sequence similarity but use random cross-validation for model selection (Partial). Per-residue attribution denotes position-level importance; DELPHI uses Integrated Gradients at raw sequence positions, whereas Chen et al. and Yu et al. report region- or feature-level importance only. Protein language models compared counts distinct pretrained models evaluated. DELPHI's training data are proprietary; its source code and trained PSR and SEC model weights are openly released (see Data and Code Availability).

| Feature | Chen et al. (1) | Yu et al. (2) | Amini et al. (3) | Abeer et al. (4) | DELPHI (this work) |
|---|---|---|---|---|---|
| Developability liabilities | Polyreactivity | Polyreactivity | Polyreactivity, HIC, self-association | SEC monomer purity | Polyreactivity, SEC monomer purity |
| Protein language models compared | 1 | 3 | 3 | 3 | 5 |
| Leakage-aware cross-validation | No | No | Yes | Partial | Yes |
| Training-set-size (learning-curve) analysis | No | No | No | No | Yes |
| Per-residue attribution | No | No | No | No | Yes |
| External validation | Independent library, clinical mAbs | 13 approved mAbs | Internal hold-out | Internal hold-out | Jain 2017, blinded Ginkgo GDPa3, public DS1 |
| Open-source software with released model weights | No | No | No | No | Yes |
